# Substrate Coupling and Inhibition of the Human Na^+^-Dependent Cl^−^/HCO₃^−^ Exchanger

**DOI:** 10.64898/2025.12.15.693380

**Authors:** Peixin Sun, Xinmin Wang, Yang Huang, Jinlun Kylian Zhang, Xurui Shen, Anqi Dong, Shiliang Wang, Xiaojun Guo, Guoming Shi, Jing Ding, Yudan Chi, Hanting Yang

## Abstract

Maintaining pH homeostasis is essential for neuronal excitability and for regulating cerebrospinal fluid (CSF). The Na^+^-dependent Cl^−^/HCO_3_^−^ exchanger, NCBE (SLC4A10), abundant in the choroid plexus, contributes to CSF pH control by mediating bicarbonate flux, yet its molecular mechanism has remained unclear. Here we report cryo-EM structures of human NCBE in multiple functional states, revealing conserved binding sites for Na^+^, Cl^−^, and HCO_3_^−^, as well as an unexpected K^+^ site indicating relaxed cation selectivity. The inhibitor DIDS binds above the substrate cavity, locking NCBE in an outward-facing conformation and defining a subfamily-selective inhibitory mechanism. Functional assays and molecular dynamics simulations validate ion coordination and inhibitor effects. Comparison of cryo-EM structures with an AlphaFold3 inward-facing model supports an elevator-like mechanism and completes the conformational cycle of Na^+^-coupled HCO_3_^−^ transporters. Together, these findings define the structural basis of NCBE and provide a framework for understanding ion coupling and pharmacological modulation within the SLC4 family.

## Introduction

Precise regulation of pH is essential for normal brain function^1,2^. Even subtle deviations in intracellular or extracellular pH can profoundly alter neuronal excitability and synaptic transmission^1,3,4^, whereas pathological acid-base disturbances, including acidosis and alkalosis, are associated with seizures^5–7^, coma^8^, and cognitive impairment^9^. The CO₂/HCO_3_^−^ buffering system constitutes the primary defence against pH fluctuations^10,11^. However, because bicarbonate (HCO_3_^−^) cannot freely diffuse across biological membranes, its movement depends on specialized membrane transporters, principally those of the SLC4 ^12,13^ and SLC26 ^14,15^ families. By coupling HCO_3_^−^ flux to ion gradients, these transporters play a central role in maintaining acid–base homeostasis across tissues.

Within the SLC4 family, the Na^+^-dependent Cl^−^/HCO_3_^−^ exchanger (NCBE, SLC4A10) is highly enriched in neurons and choroid plexus epithelium, where it drives Na^+^-dependent HCO_3_^−^ uptake to sustain synaptic activity ^16–18^ and cerebrospinal fluid (CSF) secretion ^19,20^. The physiological importance of NCBE is underscored by human genetic studies: a spontaneous deletion of exon 1 of *SLC4A10* was identified in a pair of monozygotic twins concordant for autism spectrum disorder (ASD) ^21^; a chromosomal translocation disrupting *SLC4A10* was reported in a patient with epilepsy and cognitive impairment ^22^; and ultra-rare biallelic variants cause a neurodevelopmental disorder (NDD) characterized by microcephaly, intellectual disability, epilepsy, and autism spectrum traits, with neuroimaging revealing slit ventricles and bilateral curvilinear nodular heterotopia ^23,24^. Consistent with these clinical observations, *Slc4a10*-deficient mice exhibit reduced ventricle size and altered pH-dependent neuronal excitability^17,18^. Together, these findings establish NCBE as a critical regulator of brain ion homeostasis and implicate its dysfunction in neurodevelopmental and neurological disease.

Despite its physiological and clinical importance, the structural basis of NCBE-mediated transport remains unknown. Structures of related SLC4 transporters, including AE1–3 (SLC4A1-3, Cl^−^/HCO_3_^−^ exchangers) ^25–32^, NBCe1 (SLC4A4, Na^+^-HCO_3_^−^ cotransporter) ^33^, and NDCBE (SLC4A8, Na^+^-drive Cl^−^/HCO_3_^−^ exchanger) ^34^, have revealed principles of ion coordination and conformational cycling. However, how NCBE coordinates multiple substrates, achieves ion selectivity, and is inhibited by pharmacological agents has remained unclear.

Here, we present high-resolution cryo-electron microscopy (cryo-EM) structures of human NCBE in multiple substrate- and inhibitor-bound states, complemented by molecular dynamics (MD) simulations and fluorescence-based functional assays. These structures define conserved sites for Na^+^, Cl^−^, and HCO_3_^−^, and uncover an unexpected K^+^ site that indicates relaxed ion selectivity. We further show that the stilbene derivative DIDS binds within the extracellular vestibule, occluding access to the transport pathway and stabilizing an outward-facing conformation. Comparison with AlphaFold-predicted inward-facing model supports an elevator-like mechanism of transport. These findings provide the structural framework for NCBE function and pharmacological inhibition, and broaden the mechanistic understanding of Na^+^-dependent Cl^−^/HCO_3_^−^ exchange in the brain.

### NCBE in the choroid plexus mediates HCO_3_^−^-dependent pH recovery

Single-cell transcriptomic datasets from human^35^ and mouse^36^ brain revealed that *SLC4A10*, encoding NCBE, is highly enriched in choroid plexus (CP) epithelial cells, with additional expression detected in neurons **(Fig. 1a, b; Supplementary Fig. 1a-d)**. Among SLC4 family members, NCBE showed the strongest enrichment in the human CP epithelial cells **(Supplementary Fig. 1b)**. Co-labelling with transthyretin (Ttr), a marker of choroid plexus epithelial cells (CPECs), and CD31, a vascular endothelial marker, localized NCBE to the basolateral membrane of CPECs facing the blood vessels **(Fig. 1c)**. Consistently, immunoblotting of adult mouse brain tissues confirmed robust Ncbe expression in the CP, with minimal expression detected in other regions (**Supplementary Fig. 1e**). As a HCO_3_^−^ transporter that mediates bicarbonate influx, the basolateral localization of NCBE suggests a role in vectorial ion transport and pH regulation within the CP.

**Fig. 1.**
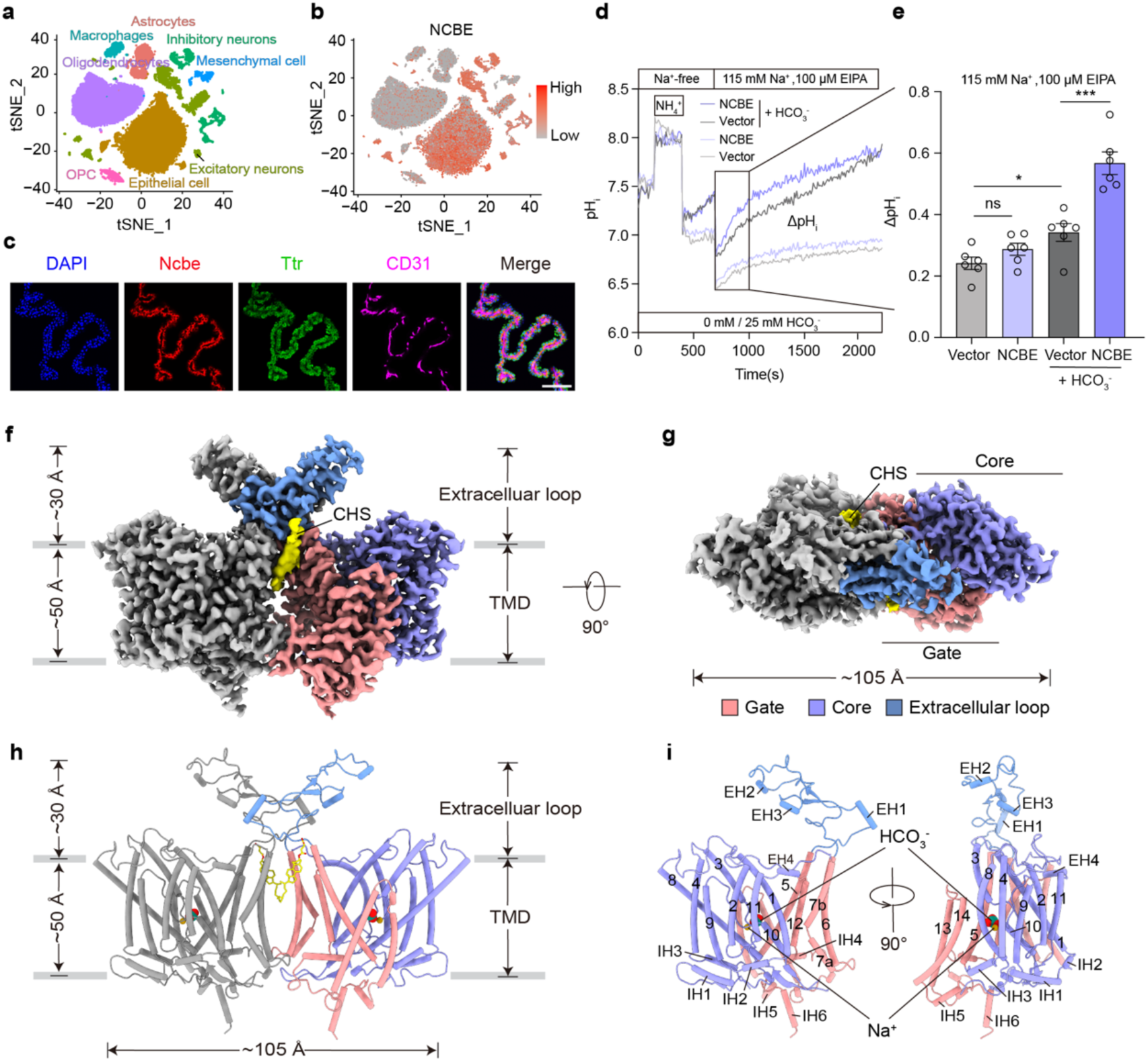
Functional characterization and cryo-EM structure of NCBE. **a, b** T-distributed stochastic neighbour embedding (T-SNE) plots reveal cell types in human medial frontal gyrus and choroid plexus samples (a) and human NCBE expression in these samples (b). **c** Immunofluorescence staining in mouse brain showing Ncbe localization in the choroid plexus. DAPI, nuclear. Ttr, a marker of choroid plexus epithelial cells. CD31, a marker of vascular endothelial cells. Scale bar 100 μm. **d,e** Time course of pH_i_ recovery (d) and statistical analysis (e) of ΔpH_i_ in NCBE-overexpressed HEK293T cells with or without HCO_3_^−^ addition. Cells transfected with vector were served as control. ΔpH_i_ was calculated as the difference between pH_i_ at 700 s and 1000 s after the addition of 115 mM Na^+^ and 100 μM EIPA. The experiments were performed in *n* = 6 biological replicates with each in technical triplicate. Data are presented as mean ± SEM. Statistical analysis was performed using unpaired t-test; \*\*\**P* < 0.001, \**P* < 0.05, ns, *P* ≥ 0.05. **f, g** Cryo-EM map of NCBE-NaHCO_3_ presented in side view (f) and top view (g). EM densities for CHS are coloured in yellow. One protomer of NCBE is coloured in grey, while the other protomer is segmented into three distinct regions: the core domain in purple, the gate domain in red and the extracellular loop in blue. **h, i** Structure representations of the NCBE-NaHCO_3_ dimer (h) and monomer, shown in cylinder mode (i). NCBE-NaHCO_3_ dimer (h) is shown from the same orientation as in (f). NCBE-NaHCO_3_ monomer (i) is shown from the side view. Each protomer of NCBE is coloured to the same scheme as shown in (f) and (g).

To directly assess HCO_3_^−^ transport activity, we monitored intracellular pH (pHᵢ) using the fluorescent dye 2′,7′-bis-(2-carboxyethyl)-5-(and-6)-carboxyfluorescein (BCECF) (**Supplementary Fig. 1f,g**)^16,37, 38^. As validation, application of NaHCO_3_ to HEK293T cells induced a progressive, time-dependent alkalinization that scaled with extracellular HCO_3_^−^ concentration, confirming that BCECF fluorescence reliably reports HCO_3_^−^-driven changes in pHᵢ **(Supplementary Fig. 1h-k)**.

We next performed a pHᵢ recovery assay in HEK293T cells expressing full-length human NCBE^16^. Following acid loading with NH₄Cl, perfusion with NaHCO_3_-containing buffer elicited a robust recovery of pHᵢ that was strictly dependent on extracellular Na^+^ **(Fig. 1d, e; Supplementary Fig. 1l, m)**. Together, these results establish NCBE as a basolateral choroid plexus transporter that mediates Na^+^-dependent HCO_3_^−^ uptake, supporting its role in CSF secretion and brain pH homeostasis.

### Cryo-EM structure reveals conserved SLC4 architecture of NCBE

To provide a framework for understanding the transport mechanism of NCBE, we purified human NCBE in the presence of NaHCO_3_ and determined its structure by single-particle cryo-EM, yielding a reconstruction at 2.45 Å resolution **(Fig. 1f-i; Supplementary Fig. 2 and Supplementary Table 1)**. NCBE assembles as a symmetric homodimer, with each protomer comprises 14 transmembrane helices (TMs) arranged into two domains, a scaffold-like gate domain (TMs 5–7 and 12–14) and a transport-active core domain (TMs 1–4 and 8–11) **(Supplementary Fig. 3a)**.

A hallmark feature of Na^+^-dependent SLC4 transporters is the extended extracellular loop between TM5 and TM6 (**Fig. 1f-i; Supplementary Fig. 3a, b**). In NCBE, this loop folds into a rigid extracellular cap stabilized by three short helices (EH1-EH3), a β-hairpin, and two disulfide bonds (C667-C715 and C669-C703) (**Supplementary Fig. 3b-d**). Both inter- and intra-protomer hydrogen-bonding interactions within this region further reinforce dimer stability **(Supplementary Fig. 3c)**. In addition, density consistent with a sterol-like molecule was observed at the extracellular dimer interface between TM5, TM6, and TM7 **(Supplementary Fig. 3e)**.

### A conserved Na^+^/HCO_3_^−^ binding pocket in NCBE

To define the substrate-binding site of NCBE, we performed detailed structural analysis of NCBE determined in the presence of NaHCO_3_. A well-defined pocket formed by TMs 3, 8, and 10 contains coordinated Na^+^ and HCO_3_^−^ density (**Fig. 2a**). The Na^+^ is coordinated by the side chain and backbone of D831, the side chain of T835, and the backbone of A876. The remaining pocket volume is occupied by density attributable to HCO_3_^−^, which is stabilized through hydrogen-bonding interactions with T835, T878, and an adjacent water molecule. Notably, HCO_3_^−^ also directly ligates the Na^+^ ion, resulting in a distorted square-pyramidal coordination geometry **(Fig. 2b, c; Supplementary Fig. 4a)**. Molecular dynamics (MD) simulations initiated from the cryo-EM model confirmed stable coordination of both Na^+^ and HCO_3_^−^ throughout the trajectories, supporting the structural assignment and the coupled nature of substrate binding **(Fig. 2d; Supplementary Table 2)**.

**Fig. 2.**
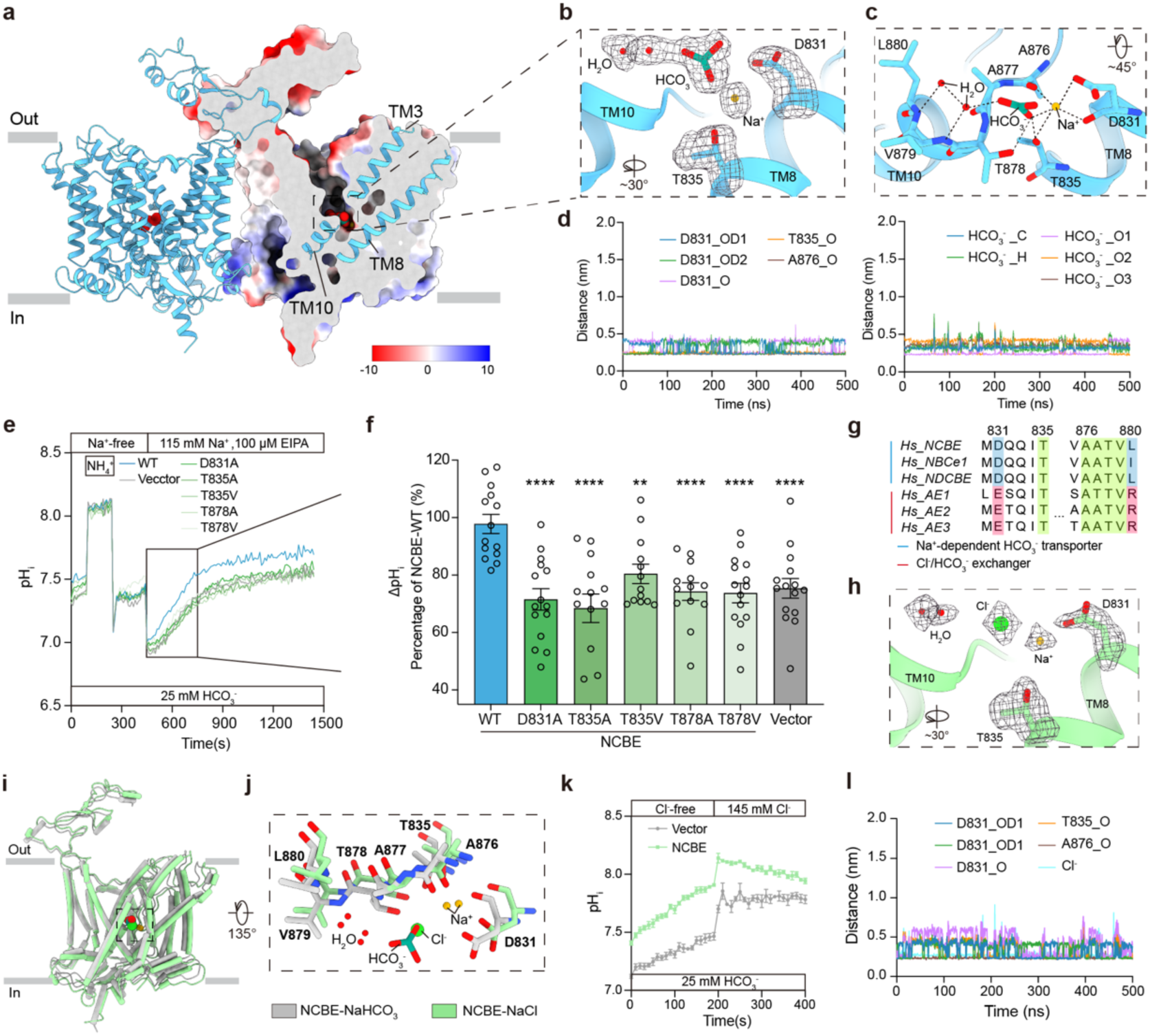
Na^+^, HCO_3_^−^ and Cl^−^ shares the conserved substrate pocket. **a** Structure and electrostatic potential surface map of NBCE NaHCO_3_-bound state. The electrostatic surface map is sliced to show the HCO_3_^−^ ion bound, with the helices TM3, TM8, and TM10 shown as blue-ribbon model. Regions with negative to positive charge are represented using a colour gradient from red to blue. **b** Zoom-in view of cryo-EM density for Na^+^, HCO_3_^−^ and nearby residues in NCBE NaHCO_3_-bound state within the conserved substrate-binding pocket. The density of ions, water and residues zoomed-in view is shown in black meshes at 0.125 level thresholds. **c** Detailed view of Na^+^ and HCO_3_^−^ binding site with key residues and coordination distances (∼ 3 Å). Residues coordinated with H_2_O, Na^+^, and HCO_3_^−^ are depicted as the atom sticks model. **d** MD simulation of 500 ns time scale illustrating the distance between the Na^+^ and its coordinated residues (left), as well as the distance between the Na^+^ and the HCO_3_^−^ (right). **e, f** Time course of pH_i_ recovery (e) and statistical analysis of ΔpH_i_ (f) in HEK293T cells overexpressing NCBE WT or its mutations. Cells transfected with the vector served as control. ΔpH_i_ was calculated as the difference between pH at 450 s and 770 s after the addition of 115 mM Na^+^ and 100 μM EIPA. The experiments were performed in *n* = 5 biological replicates with each in technical triplicate. Data are presented as mean ± SEM. Statistical analysis was performed using unpaired t-test; \*\*\*\**P* < 0.0001, \*\**P* < 0.01. **g** Sequence alignment of SLC4 family members and the positions of residues involved in binding with Na^+^ and HCO_3_^−^ are highlighted. **h** Zoom-in view of cryo-EM density for Na^+^, Cl^−^ and nearby residues in NCBE NaCl-bound state within the conserved substrate-binding pocket. The density of ions, water and residues zoomed-in view was shown in black meshes at 0.125 level thresholds. **i** The TMD superposition of NCBE NaCl-bound state (green) and NCBE-NaHCO_3_-bound state (grey) viewed from the parallel side to the cell membrane. **j** Comparison of ion-binding site from NCBE NaCl-bound state (green) and NCBE-NaHCO_3_-bound state (grey). Water molecules are shown in red ball, HCO_3_^−^ is shown in green stick, and Na^+^ and Cl^−^ are shown in yellow ball and green ball, respectively. **k** Time course of pH_i_ recovery in NCBE-overexpressed HEK293T cells with Cl^−^-free or 145 mM Cl^−^ treatment. Cells transfected with vector were served as control. The experiments were performed in *n* = 5 biological replicates with each in technical triplicate. Data are presented as mean ± SEM. **l** Distance between Na^+^, Cl^-^ and its coordinated residues (D831, T835 and A876) in 500 ns MD simulation validating stable Cl^-^ binding.

To assess the functional importance of this pocket, we expressed NCBE mutants in HEK293T cells and measured HCO_3_^−^-dependent pHᵢ recovery. Substitutions at D831, T835 and T878 markedly reduced HCO_3_^−^-dependent pHᵢ recovery relative to wild type NCBE, consistent with impaired substrate coordination. Disruption of either the Na^+^-or HCO_3_^−^-coordinating network compromised transport activity, underscoring that NCBE relies on a tightly coupled mechanism of substrate recognition **(Fig. 2e, f; Supplementary Fig. 4b)**. Sequence alignment and structural comparison across the SLC4 family revealed strong conservation of these coordinating residues among Na^+^-coupled HCO_3_^−^ transporters, including NBCe1, NDCBE, and NCBE, but not among Cl^−^/HCO_3_^−^ exchangers such as AE1–3. This divergence provides a structural basis for coupling specificity within the SLC4 family **(Fig. 2g; Supplementary Fig. 5, 6)**.

### Cl^−^ shares the conserved substrate pocket with HCO_3_^−^

NCBE has been proposed to operate either as an electroneutral Na^+^/HCO_3_^−^ cotransporter coupled to Cl^−^ self-exchange^39^ or through a Na^+^-dependent HCO_3_^−^ import-Cl^−^ export mechanism^40^; however, these models remain debated, and direct structural evidence has been lacking.

To examine the role of Cl^−^, we determined a cryo-electron microscopy structure of NCBE in the presence of NaCl at 2.6 Å resolution (**Supplementary Fig. 7, 8a; Supplementary Table 1**). The overall conformation closely resembled that of the NaHCO_3_-bound state, adopting an outward-facing architecture. Within the conserved pocket formed by TMs 3, 8, and 10, we observed well-resolved non-protein density that precisely overlapped with the HCO_3_^−^ position in the NaHCO_3_-bound structure **(Fig. 2h-j; Supplementary Fig. 8b, c)**. Under NaCl conditions, the coordination environment of this density matches the expected features of a bound anion at this site, indicating occupancy by Cl^−^.

The Na^+^ coordination network, comprising D831, T835, and the backbone carbonyl of A876, remained unchanged. The anion density was stabilized by interactions with the backbone and side chain of T878, electrostatic coupling to Na^+^, and a hydration network involving two ordered water molecules bridging the main-chain amides of T878-V879-L880 (**Supplementary Fig. 8d**). Consistent with shared occupancy of this pocket, extracellular Cl^−^ strongly inhibited Na^+^-dependent HCO_3_^−^ uptake in intracellular pH recovery assays: robust alkalinization observed in Cl^−^-free buffer was markedly suppressed upon addition of Cl^−^ (**Fig. 2k**). MD simulations further supported stable anion occupancy within this pocket over 500 ns (**Fig. 2l; Supplementary Table 3**). Collectively, these structural, functional, and computational analyses indicate that Cl^−^ and HCO_3_^−^ share a conserved binding pocket in NCBE, providing a mechanistic basis for coupling between Na^+^/HCO_3_^−^ transport and Cl^−^ binding.

### K^+^ binding at the canonical Na^+^ site reveals relaxed ion selectivity

Previous studies reported that mouse *Slc4a10* can mediate K^+^-driven Cl^−^/HCO_3_^−^ exchange^41^, suggesting that NCBE may tolerate alternative monovalent cations. To examine whether human NCBE exhibits similar behavior, we determined structure of human NCBE in the presence of KHCO_3_ at 2.5 Å resolution **(Fig. 3a; Supplementary Fig.9; Supplementary Table 1**).

**Fig. 3.**
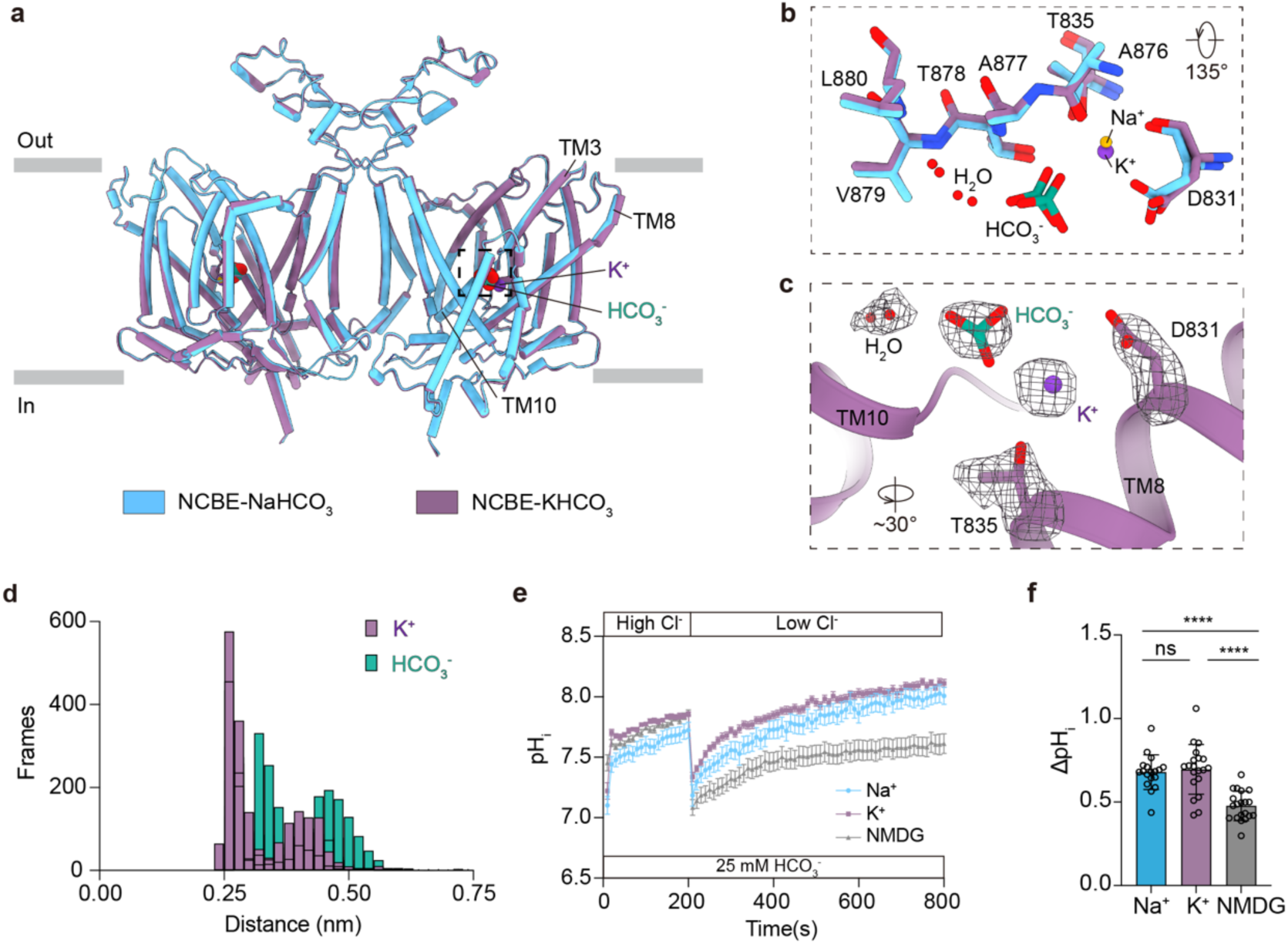
K^+^ binding at the canonical Na^+^ site reveals relaxed ion selectivity. **a** Superposition of the NCBE KHCO_3_-bound state (purple) and NCBE-NaHCO_3_-bound state (blue) from sideview. **b** Zoom-in view of cryo-EM density for K^+^, HCO_3_^−^ and nearby residues in NCBE KHCO_3_-bound state within the conserved substrate-binding pocket. The density of ions, water and residues zoomed-in view was shown in black meshes at 0.125 level thresholds. **c** Comparison of the ion-binding site from the NCBE KHCO_3_-bound state (purple) and the NCBE NaHCO_3_-bound state (blue). Water molecules are shown in red sticks, HCO_3_^−^ is shown in green sticks, and Na^+^ and K^+^ are shown in yellow ball and purple ball, respectively. **d** Histogram distribution of the distance between the K^+^ and its coordinated residues (purple), as well as the distance between the K^+^ and the HCO_3_^−^ (green). **e, f** Time course of pH_i_ recovery (e) and statistical analysis of ΔpH_i_ (f) under Na^+^-, K^+^- and NMDG^+^-containing buffer treatment in NCBE-overexpressed HEK293T cells. ΔpH_i_ was calculated as the difference between pH at 200 s and 600 s. The experiments were performed in *n* = 5 biological replicates, with each in technical triplicate. Data are presented as mean ± SEM. Statistical analysis was performed using unpaired t-test; \*\*\*\**P* < 0.0001, ns, *P* ≥ 0.05.

Structural superposition of the KHCO_3_- and NaHCO_3_-bound states revealed only minor backbone deviations within the substrate-binding pocket. In the KHCO_3_-bound structure, the canonical Na^+^ site between TMs 3, 8, and 10 is occupied by a cation density that closely overlaps with the Na^+^ position observed in the NaHCO_3_-bound structure, while density corresponding to HCO_3_^−^ and two ordered water molecules remains preserved **(Fig. 3a-c; Supplementary Fig. 10a, b)**. Under KHCO_3_ conditions, the coordination features at this site are compatible with K^+^ binding, indicating that the pocket can accommodate an alternative monovalent cation without substantial rearrangement.

All-atom MD simulations simulations performed in a K^+^-only cation environment further supported stable occupancy of this site by K^+^, with coordination involving D831, T835, and the backbone carbonyl of A876, while maintaining electrostatic association of HCO_3_^−^ **(Fig. 3d; Supplementary Fig. 10c, d; Supplementary Table 4)**. Intracellular pH recovery assays further revealed pronounced alkalinization in the presence of either Na^+^- or K^+^-containing buffers, whereas no significant alkalinization was observed when both cations were replaced by NMDG **(Fig. 3e, f)**. These results indicate that the canonical Na^+^ site in NCBE exhibits relaxed selectivity toward monovalent cations, revealing intrinsic flexibility of the substrate-binding pocket in the human transporter.

### Elevator-like conformational transition of NCBE

Given an inward-facing (IF) conformation of NCBE has not been resolved by cryo-electron microscopy, we used AlphaFold3 (AF3) ^42^ to predict this missing state **(Fig. 4a, left; Fig. 4b)**. To place the experimentally determined outward-facing (OF) conformations into a mechanistic framework, we compared our cryo-EM structures with the AF3-predicted IF model. The AF3 model revealed a deep cytoplasmic vestibule, whereas all three ion-bound cryo-EM structures, captured in NaHCO_3_, KHCO_3_, and NaCl, adopted outward-facing conformations characterized by an open extracellular vestibule **(Fig. 4a-c)**.

**Fig. 4.**
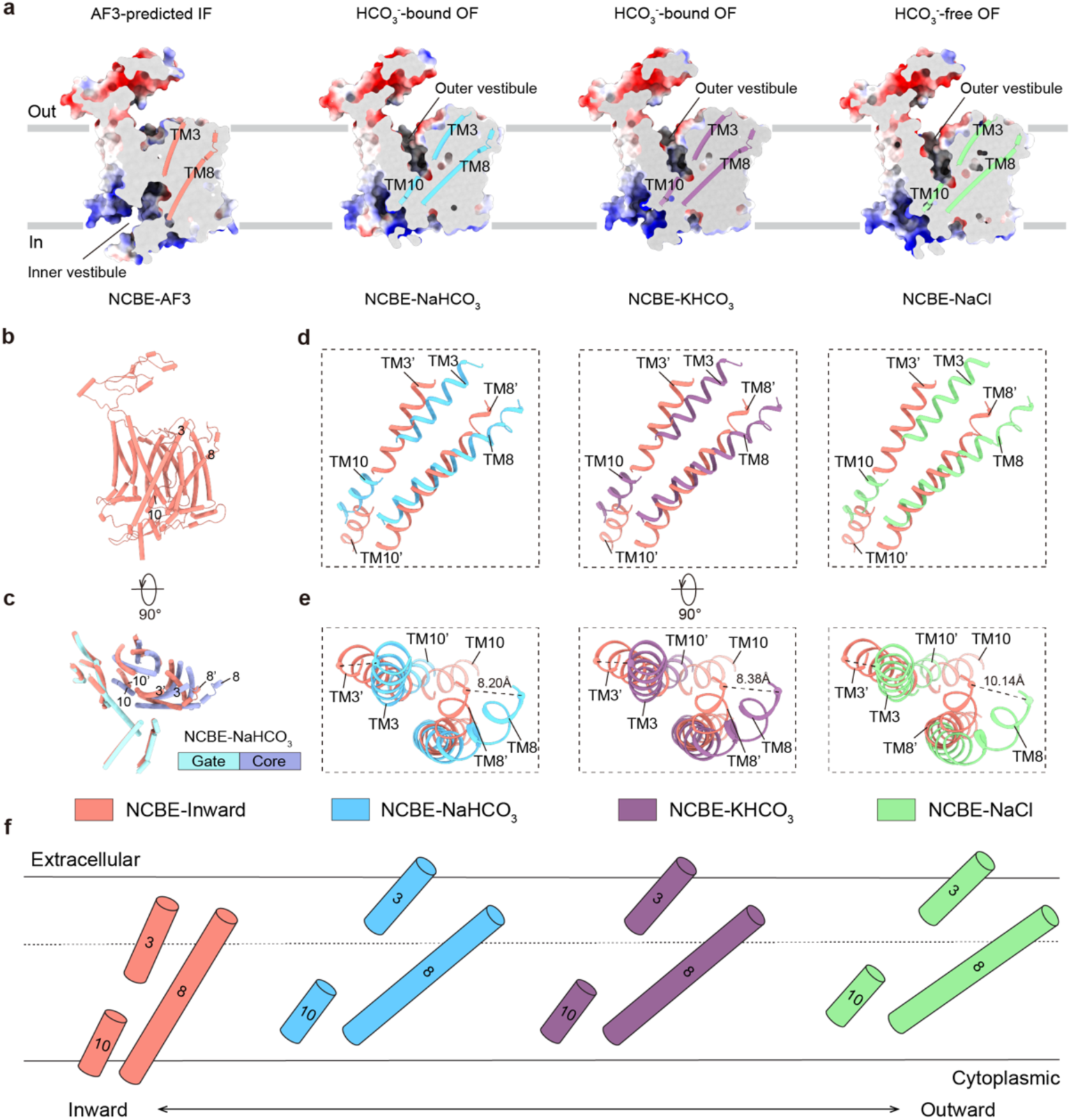
Conformational transitions of NCBE between IF and OF states. **a** Electrostatic potential surface map of monomer NBCE with different substrates. The electrostatic surface map is sliced to show the vestibule, with the helices TM3, TM8, and TM10 shown as cylinder models. The TM3, TM8, and TM10 of NCBE-AF3, NCBE-NaHCO_3_, NCBE-KHCO_3_ and NCBE-NaCl are coloured by light coral, blue, purple, and green, respectively. Regions with negative to positive charge are represented using a colour gradient from red to blue. **b** AF3-predicted IF state of NCBE structure. **c** Structural alignment of NaHCO_3_-bound and the AF3-predicted NCBE structure. The gate domain was used as the reference for structural alignment. Substrate-binding helices in transmembrane segments TM3, TM8, and TM10 are labelled. **d, e** Different views of the schematic depict conformational differences between NCBE with different substrate states. TM3, TM8, and TM10 are shown in ribbon model. **f** Schematic representation of the conformational differences between the OF state of NCBE in various substrate-bound and the AF3-predicted IF state of NCBE. Transmembrane helices TM3, TM8, and TM10 are depicted using cartoon models.

Comparison of the AF3-predicted IF model with the cryo-EM OF structures indicates a pronounced vertical displacement of the transport-active core domain relative to the scaffold-like gate domain. This movement is driven primarily by concerted rearrangements of TMs 3, 8, and 10 and spans approximately 8–10 Å between the IF and OF states (**Fig. 4d, e**). Both the magnitude and direction of the displacement were conserved across all ion-bound structures, indicating that different substrates stabilize outward-facing conformations without altering the fundamental transport trajectory.

To illustrate this conformational transition, we generated a schematic model of the transport cycle **(Fig. 4f)**. In this model, the core domain behaves as a rigid body that alternates between inward- and outward-facing states, carrying bound Na^+^, K^+^, HCO_3_^−^, and Cl^−^ across the membrane via coordinated movement of TMs 3, 8, and 10, while the gate domain remains comparatively stationary. This mode of motion is characteristic of an elevator-type mechanism, distinguishing NCBE from the rocking-bundle or rocker-switch mechanisms described for other classes of secondary transporters ^43, 44^.

The observed conformational differences support an elevator-like conformational transition in NCBE, providing a mechanistic framework for alternating access in a Na^+^/K^+^-dependent Cl^−^/HCO_3_^−^ exchanger and extending current models within the SLC4 family.

### DIDS occludes the extracellular vestibule and stabilizes the outward-facing state

DIDS (4,4′-diisothiocyanatostilbene-2,2′-disulfonic acid) is a classical stilbene-based inhibitor of bicarbonate transporters ^12^. In HEK293T cells expressing NCBE, DIDS treatment markedly suppressed HCO_3_^−^ uptake, as evidenced by a strong reduction in intracellular pH recovery following acid loading **(Fig. 5a, b)**.

**Fig. 5.**
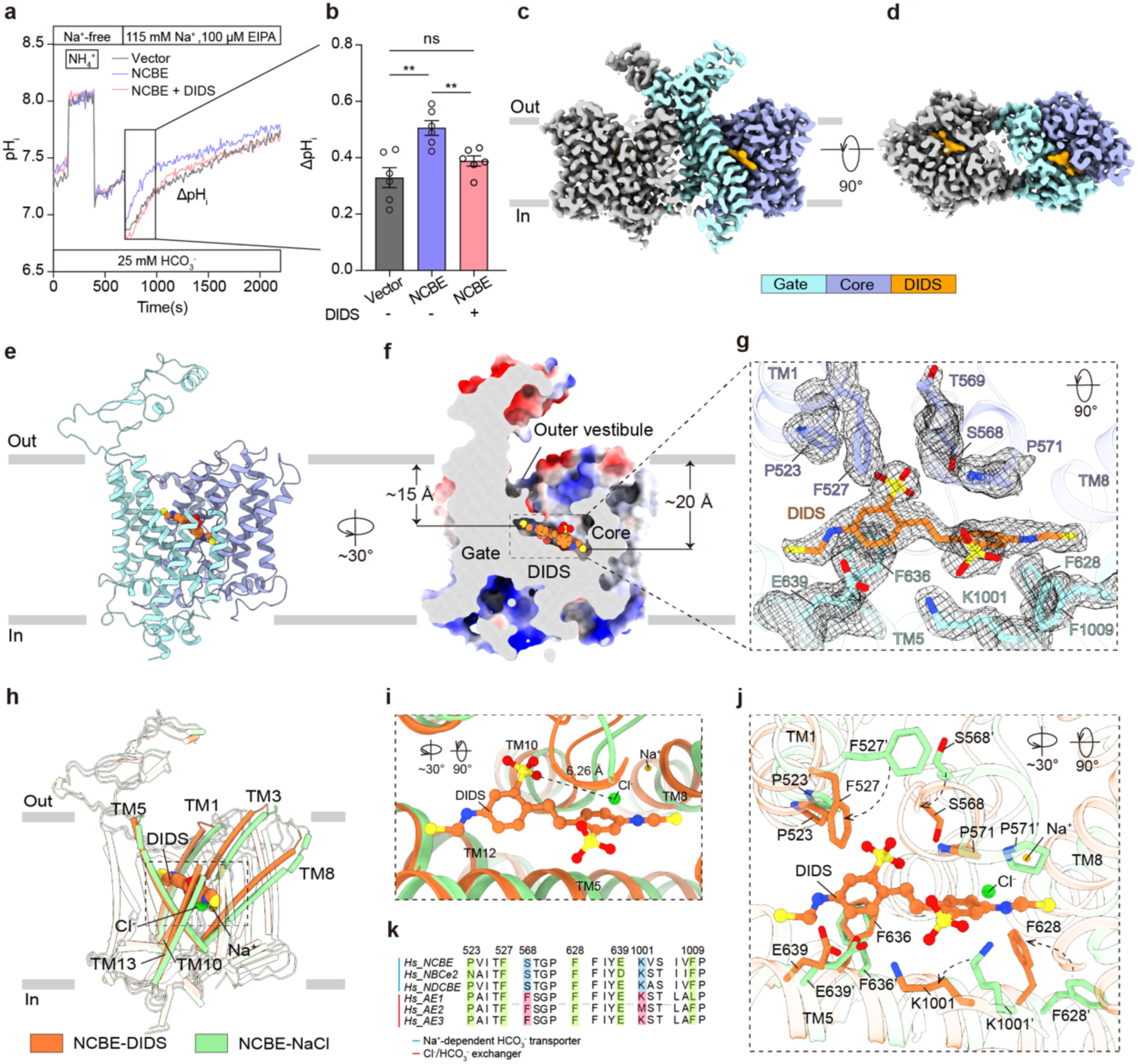
DIDS binding occludes the extracellular vestibule of NCBE. **a, b** Time course of HCO_3_^−^-dependent pH recovery (a) and statistical analysis of ΔpH_i_ (b) in NCBE-overexpressed HEK293T cells without or with 300 μM DIDS treatment. Cells transfected with the vector served as a control. ΔpH_i_ was calculated as the difference between pH at 700 s and 1000 s after the addition of 115 mM Na^-^ and 100 μM EIPA. The experiments were performed in *n* = 6 biological replicates with each in technical triplicate. Data are presented as mean ± SEM. Statistical analysis was performed using unpaired t-test; ** *P* < 0.01, ns *P* > 0.05. **c, d** Cryo-EM density map of NCBE-DIDS is sliced to show the DIDS and presented in two different views: side view (c) and top view (d). Electron microscopy densities for DIDS are coloured in orange. One protomer of NCBE is coloured in grey, while the other protomer is segmented into two distinct regions: the core domain in purple, the gate domain in cyan. **e, f** Ribbon model (e) and electrostatic potential surface map (f) of NBCE-DIDS monomer structure. The electrostatic potential surface map is sliced to show the DIDS sphere model in the extracellular cleft. DIDS is shown as orange spheres. **g** Cryo-EM density for DIDS and nearby residues in NCBE-DIDS state within the binding pocket. The density of DIDS and residues zoomed-in view was shown in black meshes at 0.176 level thresholds. **h** Comparison of NCBE-DIDS (orange) and NCBE-NaCl (green) structures. Critical TMs are represented by coloured cylinder without transparency. DIDS, Na^+^ and Cl^-^ are shown as sphere model. **i** Comparison of DIDS binding pocket and NaCl binding pocket. DIDS, Na^+^ and Cl^-^ are shown as stick model. **j** Zoom-in view of the binding pock and interactions of the DIDS in the NCBE-DIDS (orange) and NCBE-NaCl (green) structures. The DIDS is shown as orange stick model. Na^+^ and Cl^-^are shown as yellow and green stick model. Critical residues interacting with the ligands are shown as sticks and labelled. **k** Sequence alignment of SLC4 family members and the positions of residues involved in binding with DIDS are highlighted.

In the DIDS-bound cryo-EM structure, NCBE adopts an outward-facing conformation, with the inhibitor lodged within the extracellular vestibule between the core and gate domains (**Fig. 5c, d**). DIDS is oriented implicitly relative to the membrane plane, extending from the extracellular surface toward the substrate-binding cavity, with its two isothiocyanate groups projecting approximately 15 Å and 20 Å, respectively (**Fig. 5e, f**). DIDS is anchored by hydrophobic interactions with P523, F527, F628, F636, and F1009, together with polar contacts involving S568 and K1001 (**Fig. 5g**). Occupation of the vestibule blocks the access pathway to the ion-binding pocket, sterically preventing substrate entry and constraining the conformational transitions required for transport (**Fig. 5h, i; Supplementary Fig. 12**). One sulfonic acid group of DIDS lies within ∼3-6 Å of the HCO_3_^−^/Cl^−^ binding site, suggesting potential electrostatic interference with anion binding. Local rearrangements of residues including F527, S568, P571, F628, F636, and K1001 further accommodate and stabilize the bound inhibitor (**Fig. 5j; Supplementary Fig. 12c, f**). MD simulations confirmed persistent occupancy of DIDS within this vestibule over 500 ns (**Supplementary Table 5**).

In contrast to AE1-3, where DIDS forms covalent adducts, the inhibitor in NCBE resides ∼6 Å deeper in the extracellular vestibule and is positioned ∼6.4 Å from K643, precluding covalent linkage. Instead, stabilization is mediated by a hydrogen bond with S568, a residue replaced by phenylalanine in Cl^−^/HCO_3_^−^ exchangers (**Fig. 5k; Supplementary Fig. 13**), highlighting a distinct inhibitory mechanism for Na^+^-coupled bicarbonate transporters.

## Discussion

NCBE (SLC4A10) enables Na^+^-dependent bicarbonate transport in the brain by integrating multi-ion recognition with large-scale conformational rearrangements. The structures presented here reveal how a conserved substrate pocket accommodates Na^+^, HCO_3_^−^, Cl^−^, and K^+^, how vertical movement of the transport core provides alternating access, and how pharmacological inhibition is achieved by extracellular occlusion. Together, these features define the molecular operating principles of a Na^+^-dependent Cl^−^/HCO_3_^−^ exchanger (**Supplementary Fig. 15**).

NCBE is enriched at the basolateral membrane of choroid plexus epithelial cells, where it contributes to cerebrospinal fluid secretion^45^ and pH regulation^20^. The Na^+^-dependent HCO_3_^−^ influx observed here offers a structural explanation for phenotypes associated with *Slc4a10* deficiency, including altered CSF composition and seizure susceptibility^18^, and provides a framework for interpreting neurological disorders linked to *SLC4A10* variants.

At the core of NCBE function is a conserved ion-binding pocket that couples cation and anion recognition. Mutations disrupting Na^+^/HCO_3_^−^ coordination impair bicarbonate-driven alkalinization (**Fig. 2e, f**), underscoring tight substrate coupling. The ability of this pocket to also accommodate K^+^ (**Fig. 3a-d**), in line with prior observations in mouse *Slc4a10*^41^, indicates relaxed monovalent cation selectivity relative to other Na^+^-dependent SLC4 transporters.

Cl^−^ interaction with NCBE has been controversial, with models proposing either self-exchange^39^ or net export coupled to bicarbonate uptake^40^. Structural occupancy of Cl^−^ at the anion site (**Fig. 2h-j**), together with functional suppression of HCO_3_^−^ uptake by extracellular Cl^−^ (**Fig. 2k**), supports direct competition between anions at a shared pocket, reconciling these models while preserving a common SLC4 architecture.

Across multiple substrate and inhibitor conditions, NCBE adopts outward-facing conformations with an open extracellular vestibule (**Fig. 4a-c**). Comparison with an AlphaFold3-predicted inward-facing state indicates a vertical translation of the transport core relative to the gate domain **(∼8-10 Å; Fig. 4d-f**), consistent with an elevator-type mechanism. Similar motions observed in other SLC4 transporters^25–34^ suggest that vertical core displacement is a conserved strategy for coordinating multiple ions at a single site.

Inhibition by DIDS occurs through occupation of the extracellular vestibule, stabilizing an outward-facing state and blocking substrate access (**Fig. 5c–i**). Unlike Cl^−^/HCO_3_^−^ exchangers, where DIDS forms covalent adducts, NCBE binds the inhibitor through non-covalent interactions involving S568 (**Fig. 5k; Supplementary Fig. 13**), defining a subfamily-selective inhibitory mode.

Overall, by resolving how Na^+^, K^+^, Cl^−^, and HCO_3_^−^ converge on a single conserved pocket, how elevator-like core movements enable alternating access, and how DIDS exploits the extracellular vestibule to arrest transport, this study establishes a mechanistic blueprint for NCBE function. These insights extend fundamental principles of SLC4 transport in the brain and offer a structural foundation for interpreting disease-associated mutations and developing subfamily-selective modulators of bicarbonate transport.

## Methods

### Experimental animals

All animal experiments were approved by the Institutional Animal Care and Use Committee at Fudan University. C57BL/6J mice were purchased from GemPharmatech Co.. For experiments, mice aged 6 weeks were utilized and maintained in a specific pathogen-free (SPF) animal facility. They were provided with sterilized water and standard chow ad libitum and housed individually in ventilated cages under controlled environmental conditions (12-hour light/dark cycle, 22°C, 40%–60% relative humidity).

### Immunohistochemistry

After anesthesia, the mice were perfused with PBS and the brain tissues were removed. The brain tissues were fixed with 4% paraformaldehyde and dehydrated for 48 hours. Then, the brain tissues were embedded in OCT. embedding medium (4583) and rapidly frozen on dry ice. The OCT-embedded brain tissues were sectioned into 10 μm -thick slices for subsequent staining. Brain slices were incubated in blocking/permeabilization solution (PBS containing 10% donkey serum, 2% BSA and 0.4% Triton X-100) for 1 hour. Then, the slices were incubated with primary antibodies at 4°C overnight. Subsequently, they were incubated with fluorescently labelled secondary antibodies in PBS containing 1% BSA and 0.04% Triton X-100 at room temperature for 1.5 hours. Nuclei were stained with DAPI (Sigma, #D9542) for 3 minutes. Finally, the slices were mounted with Fluoro-Gel mounting medium (Electron Microscopy Sciences, #17985) and imaged using an FV3000 confocal microscope (Olympus). The following primary antibodies were used in immunofluorescence staining: anti-CD31 (550274, BD Biosciences, 1:500), anti-Transthyretin (PA5-20742, Thermo Fisher, 1:500), anti-Slc4a10 (Proteintech, 27197-1-AP, 1: 500).

### Immunoblotting

Tissue samples were collected from mice for Western blot analysis. Following anesthesia, animals underwent transcardial perfusion, and tissues were promptly excised. Harvested tissues were homogenized in RIPA lysis buffer supplemented with protease inhibitor cocktail and subjected to five cycles of ultrasonication on ice to ensure complete protein extraction. Lysates were then centrifuged at 12,000 × g for 10 min at 4°C, and supernatants containing total protein were collected. Protein concentration was quantified using a BCA Protein Assay Kit (B5001, LabLead). Equal protein amounts (20 μg per sample) were separated by SDS-PAGE and transferred onto nitrocellulose (NC) membranes. Membranes were blocked with 5% milk in TBST and incubated overnight at 4°C with the primary antibody. After three TBST washes, membranes were incubated with HRP-conjugated secondary antibodies for 1 h at room temperature. Immunoreactive bands were detected using an ECL Detection Kit (Merck Millipore, #WBKLS0500) and visualized with a Bio-Rad imaging system. The following primary antibody were used in western blotting: anti-Slc4a10 (Proteintech, 27197-1-AP, 1: 2000) and anti-GAPDH (Proteintech, 60004-1-Ig, 1: 10,000).

HEK293T cells were lysed in RIPA buffer supplemented with 1 mM PMSF for 10 min. Lysates were centrifuged at 12,000 × g for 10 min at 4 °C, and the supernatants were collected. Protein concentrations were measured using a BCA Protein Assay Kit (Beyotime, P0010). Equal amounts of protein were separated by SDS–PAGE and transferred to PVDF membranes (Cytiva) for western blot analysis. Human NCBE and its mutants were detected with a mouse anti-Flag mAb (Proteintech, 66008-4-Ig, 1:10,000), and actin was detected with a mouse anti-*Beta* actin mAb (Proteintech, 66009-1-Ig, 1:10,000) as a loading control. An HRP-conjugated goat anti-mouse IgG(H+L) (Proteintech, SA00001-1, 1:10,000) was used as the secondary antibody. All antibodies were diluted 1:10,000 in TBST containing 5% skim milk before use. Signals were visualized using a Bio-Rad imaging system.

### Full-length human NCBE expression and purification

Gene encoding the full-length human NCBE (UniprotKB-Q6U841) was cloned into the modified pCAG with a Flag tag plus an HA tag and a TEV protease site at the C terminus. The reconstructed plasmid of human NCBE was transfected into GnTI^-^ or Expi293 cells using PEI MW40000 (Yeasen), when cell density reached ∼ 2.5-3.0 × 10^6^ cells ml^-1^. The transfected GnTI^-^ or Expi293 cells were further cultured for 60 h or 72 h, respectively, and harvested by centrifugation at 1,500 × g for 10 min. Cell pellets were washed with PBS buffer three times, resuspended, and homogenized with lysis buffer (20 mM HEPES, 150 mM NaCl, pH 7.4) containing 2 μg ml^-1^ protease inhibitor cocktail (Aprotinin:Pepstatin:Leupeptin (w/w) = 1:1:1), and 1 mM PMSF. The cell membrane protein was extracted by adding 1% (w/v) n-dodecyl-β-D-maltopyranoside (DDM, Anatrace) and 0.1 % (w/v) cholesteryl hemisuccinate (CHS, Anatrace) and stirring gently at 4 °C for 2 h. The solution was centrifuged at 202,000 × g at 4 °C for 1 h. After centrifugation, the supernatant was collected for incubation with anti-Flag resin (GeneScript) at 4 °C for 1 h, and the resin was washed with lysis buffer with 0.05% (w/v) glycodiosgenin (GDN, Anatrace). The protein was eluted using the lysis buffer with 0.05% GDN and 0.5 mg ml^-1^ 3 × flag peptide. Superose 6 Increase 10/300 GL column (Cytiva) was used in size-exclusion chromatography. The mobile phase in the column contains 20 mM HEPES, 100 mM NaCl, 50 mM NaHCO_3_, pH 7.4, and 0.01% (w/v) GDN. The corresponding NCBE fractions were collected and concentrated until the protein concentration reached about 7 mg ml^-1^ for cryo-EM grids preparation. When preparing NCBE-KHCO_3_ sample, the NaCl and NaHCO_3_ were replaced by KCl and KHCO_3_, respectively, during the protein purification. When preparing NCBE-DIDS sample, the NCBE-NaCl protein sample was incubated with 0.5 mg ml^-1^ DIDS (Sigma) for 30 min at 4°C for the cryo-EM sample.

### Cryo-EM sample preparation and data collection

Aliquots of 3 μl (∼7 mg ml^-1^) of purified NCBE samples were applied to glow-discharged holey carbon grids (Quantifoil R1.2/1.3, Au, 300 mesh). The cryo-EM grids were prepared using the Leica EM-GP2 automatic plunge freezer. The whole process of cryo-EM grids preparation was carried out at 4 °C with 80% humidity. The protein sample was waiting for 10s after being added onto the grids and then blotted with filter paper for 3s-4s. The protein-coated grids were then plunged into liquid ethane and stored in liquid nitrogen for further screening. The high-resolution images were collected on Titan Krios G4 cryo-electron microscope operated at 300 kV, equipped with a Falcon 4i Direct Electron Detector and a Selectris X energy filter (Thermo Fisher Scientific). The movies were automatically collected using EPU software (Thermo Fisher) with a nominal magnification of 130,000×, which yields a final pixel size of 0.932 Å, defocus range between -0.8 and -1.8 μm. Movies were collected at 3.51 s of total exposure, corresponding to an electron dose of 49∼50 e^−^ Å^−2^.

### Data processing

The overall workflow for image processing of the NCBE-NaCl, NCBE-NaHCO_3_, NCBE-KHCO_3_, and NCBE-DIDS datasets is illustrated, and additional details can be found in the Supplementary Information. The data processing method was referred to prior research ^46^. All image processing was performed by using CryoSPARC (v.4.5.3)^47^.

For the dataset of the NCBE-NaHCO_3_, 12,649 movies were initially aligned using Patch Motion Correction and contrast transfer function (CTF) was determined using patch CTF determination. 3,943,966 particles were auto-picked using the blob picker and template picker and particles were extracted with a box size of 640 pixels and fourier crop to a box size of 320 pixels, after 2 rounds of 2D classification, 1,993,997 particles were used for ab initio reconstruction in five classes. After additional particle clean-up by further heterogeneous refinement, 1,148,840 particle sets were re-extracted with a box size of 320 pixels at pixel size of 0.932 Å and used to perform homogeneous refinement and non-uniform refinement, yielding a resolution of 2.57 Å. The final improved reconstruction map was refined to 2.45 Å from a total of 247,466 particles after local CTF refinement and non-uniform refinement with mask.

For the NCBE-KHCO_3_ dataset, 5,679 movies were first aligned with Patch Motion Correction, and CTF parameters were determined using Patch CTF Determination. A total of 1,282,181 particles were picked with the blob and template pickers and extracted with a 640-pixel box size, followed by Fourier cropping to 320 pixels. After two rounds of 2D classification, 541,673 particles were selected for ab initio reconstruction into five classes. Additional dataset cleaning through heterogeneous refinement reduced the set to 271,912 particles, which were re-extracted with a 320-pixel box size at a pixel size of 0.932 Å. These particles were refined using both homogeneous and non-uniform refinement, yielding a reconstruction at 2.61 Å resolution. After local CTF refinement and masked non-uniform refinement, the final reconstruction reached 2.51 Å from 221,965 particles.

For the NCBE-NaCl dataset, a total of 11,862 movies were initially aligned using Patch Motion Correction, and the contrast transfer function (CTF) parameters were estimated with Patch CTF Determination. Particle picking was performed with both the blob picker and template picker, yielding 2,147,632 particles that were extracted with a box size of 640 pixels and Fourier-cropped to 320 pixels. Following two rounds of 2D classification, 831,923 particles were retained for ab initio reconstruction into five classes. Subsequent heterogeneous refinement further cleaned the dataset, resulting in 544,749 particles, which were re-extracted with a 320-pixel box size at a pixel size of 0.932 Å. These particles were subjected to homogeneous refinement and non-uniform refinement, producing a reconstruction at 2.74 Å resolution. After local CTF refinement and masked non-uniform refinement, the final map was obtained at 2.62 Å resolution from 269,507 particles.

For the NCBE-DIDS dataset, 4,747 movies were processed by Patch Motion Correction, and CTF parameters were assessed through Patch CTF Determination. Particle picking with the blob and template pickers yielded 462,825 particles, which were extracted using a 640-pixel box size and Fourier-cropped to 320 pixels. Following two rounds of 2D classification, 178,679 particles were used for ab initio reconstruction into five classes. Subsequent heterogeneous refinement further reduced the dataset to 123,866 particles, which were re-extracted with a 320-pixel box size at a pixel size of 0.932 Å. These were refined through homogeneous and non-uniform refinement, producing a reconstruction at 2.75 Å resolution. With local CTF refinement and masked non-uniform refinement, the final reconstruction was improved to 2.61 Å resolution, based on 123,866 particles.

The reported resolutions are all based on the gold standard Fourier Shell Correlation (FSC) 0.143 criterion.

### Model building and refinement

An initial model of human NCBE was generated by AlphaFold3 ^42^. AF3 model was fitted into density maps using UCSF ChimeraX ^48^. The model was then manually adjusted using Coot ^49^. The local areas of the models were refined using real-space refinement with Ramachandran and geometry constraints. The full models of NCBE were then automated refined in PHENIX by using Phenix.real_space_refine ^50^. Figures were prepared using ChimeraX.

### Molecular dynamics simulations

Molecular dynamics (MD) simulation was conducted for NCBE-NaCl, NCBE-NaHCO_3_, NCBE-KHCO_3_ and NCBE-DIDS systems. MD simulation was referred to prior research ^46^. The structures and force fields for the MD system were generated utilizing the CHARMM-GUI (v.3.7) ^51^.

The CHARMM36 force field was employed for proteins, ions, POPC lipids ^52^, and TIP3P water molecules ^53^. The force field of DIDS was generated using the CHARMM General Force Field (CGenFF) program ^54,55^. Charge neutralization of NCBE-NaCl, NCBE-NaHCO_3_ and NCBE-DIDS systems was achieved through targeted addition of Na^+^, Cl^−^, and HCO_3_^−^ ions, yielding final concentrations of 0.15 M NaCl, 0.14 M NaCl and 0.01 M NaHCO_3_, and 0.15 M NaCl, respectively. The NCBE-KHCO_3_ system was neutralized with 0.14 M KCl and 0.01 M KHCO_3_. Comprehensive system compositions are documented in Supplementary Table S2-S5.

Subsequent to system preparation, energy minimization, equilibration, and MD production were conducted using GROMACS (v.2023.3) ^56^. Energy minimization employed the steepest descent algorithm, continuing until convergence was achieved (maximum force < 1000 kJ mol^−^¹ nm^−^¹) to remove steric clashes and geometric artifacts. During the equilibration phase, lasting approximately 2 ns, the system temperature was maintained at 303.15 K using the Berendsen thermostat ^57^ under NVT conditions for an initial 500 ps. This was followed by pressure regulation at 1 bar using the same thermostat under NPT conditions for an additional 1 ns. Throughout equilibration, hydrogen bond constraints were applied using the LINCS algorithm ^58^.

Following pre-equilibration, 500 ns MD production was performed. Trajectory coordinates were recorded at 10 ps intervals. System temperature was maintained at 303.15 K using the Nosé-Hoover thermostat ^59^, while pressure was regulated at 1 bar via the Parrinello-Rahman barostat ^60^. Long-range electrostatic interactions were computed with the Particle Mesh Ewald (PME) method ^61^ using a 12 Å cutoff. Van der Waals interactions were smoothly truncated between 10–12 Å using a switching function. Periodic boundary conditions (PBC) were applied throughout all simulations. The distances between substrates and coordinating residues in 500 ns MD production were calculated using the GROMACS *gmx distance* utility.

### Intracellular pH (pH_i_) calibration

HEK 293T cells were loaded with the intracellular pH indicator BCECF-AM (Thermo) at 1 μM in PBS for 1 h at 37 °C. The intracellular pH calibration was performed by using pH_i_ stepwise from pH 8.0 to 6.0 in high-K^+^ buffer (10 mM HEPES, 115 mM KCl, 1mM CaCl_2_, 1mM MgCl_2_, 0.2 mM / 0.8 mM KH_2_PO_4_/K_2_HPO_4_) with 10 μM nigericin and valinomycin mixture to treat HEK 293T cells ^40^. The BCECF fluorescence was monitored by a dual-excitation wavelength of 490 nm / 440 nm excitation and 535 nm emission by using microplate reader (Spark, TECAN). Data were analyzed by GraphPad Prism software.

### HCO_3_^−^-dependent pH recovery assay

Following the loading of BCECF-AM as outlined above, HEK 293T cells were treated with DMEM (10% FBS + 1% PS) with 0 mM, 3 mM, 10 mM, 15 mM and 25 mM NaHCO_3_ for 10 minutes. The BCECF fluorescence was monitored as outlined above. Data were analyzed by GraphPad Prism software.

### Na^+^-dependent HCO_3_^−^ transport assay

The Na^+^-dependent HCO_3_^−^ transport assay was conducted as previously described ^16^. HEK 293T cells were transfected with expression vectors encoding NCBE and various mutants for 48 hours. Following the loading of BCECF-AM as outlined above, the transfected cells were acidified using a Na^+^-free buffer containing 40 mM NH₄^+^ (10 mM HEPES, 75 mM TMA-Cl, 1 mM CaCl₂, 1 mM MgCl₂, 25 mM KHCO_3_, 40 mM NH₄Cl, 0.2 mM / 0.8 mM KH₂PO₄/K₂HPO₄, pH 7.4) for 3 minutes. To assess Na^+^-dependent HCO_3_^−^ transport, the cells were first switched to a Na^+^-free buffer (10 mM HEPES, 115 mM TMA-Cl, 1 mM CaCl₂, 1 mM MgCl₂, 25 mM KHCO_3_, 0.2 mM / 0.8 mM KH₂PO₄/K₂HPO₄, pH 7.4) for 5 minutes, and then transferred to a Na^+^-containing buffer (10 mM HEPES, 90 mM NaCl, 25 mM KCl, 1 mM CaCl₂, 1 mM MgCl₂, 25 mM NaHCO_3_, 0.2 mM / 0.8 mM KH₂PO₄/K₂HPO₄, pH 7.4). The BCECF fluorescence was monitored as previously described. Data were analyzed by GraphPad Prism software.

### Cl^−^-HCO_3_^−^ exchange activity assay

The Cl^−^-HCO_3_^−^ exchange activity assay was performed as previously described ^30^. After transfected with NCBE and loaded with BCECF-AM as mentioned above, the cells were treated by Cl^−^-free buffer (140 mM Na-Gluconate, 5 mM K-Gluconate, 1 mM MgSO_4_, 1mM Ca-Gluconate, 5 mM Glucose, 25 mM NaHCO_3_, 10 mM HEPES, 2.5 mM NaH₂PO_4_, pH 7.4) for 3 min and then transferred to 140 mM Cl^−^ buffer (140 mM NaCl, 5 mM KCl, 1 mM MgSO_4_, 1mM Ca-Gluconate, 5 mM Glucose, 25 mM NaHCO_3_, 10 mM HEPES, 2.5 mM NaH₂PO_4_, pH 7.4). The BCECF fluorescence was monitored as previously described. Data were analyzed by GraphPad Prism software.

### K^+^-dependent HCO_3_^−^ transport assay

The K^+^-dependent HCO_3_^−^ transport assay was performed as previously described ^41^. After transfection with NCBE and loading with BCECF-AM as described above, cells were incubated in high Cl^−^ buffer (10 mM HEPES, 5 mM glucose, 25 mM NaHCO_3_, 120 mM KCl, 1 mM CaCl₂, 1 mM MgCl₂, pH 7.4) for 3 min, followed by incubation in low Cl^−^ buffer (10 mM HEPES, 5 mM glucose, 25 mM KHCO_3_, 120 mM K-gluconate, 1 mM CaCl₂, 1 mM MgCl₂, pH 7.4) for 10 min. Cells treated with Na^+^- or NMDG-based buffer were used as controls. In the NMDG buffer, NaHCO_3_ or KHCO_3_ was replaced by choline-HCO_3_. BCECF fluorescence was recorded as previously described, and data were analyzed using GraphPad Prism.

## Statistical information

All data were analyzed using GraphPad Prism software (version 9.5.1). For comparison of two groups, unpaired Student’s t test was used. Data in bar graphs are presented as mean ± SEM. Levels of statistical significance are indicated as follows: *p <0.05, **p <0.01, ***p < 0.001, and **** p < 0.0001.

## Data and code availability

The GEO accession ID of scRNA-seq from human and mouse are GSE159812 and GSE168704. The Protein Data Bank coordinates used for structural alignments are 7TY7 (human AE1 with HCO_3_^−^), 8GVC (human AE2 with HCO_3_^−^), 8Y86 (human AE3 with HCO_3_^−^), 6CAA (human NBCe1), 7RTM (European rabbit NDCBE with NaHCO_3_), 8T6V (human AE1 with DIDS), 8GV8(human AE2 with DIDS) and 8Y85(human AE3 with DIDS). The UniProt accession code of human SLC4A10 (NCBE) is Q6U841. The 3D cryo-EM density maps of human NCBE-NaHCO_3_, NCBE-KHCO_3_, NCBE-NaCl and NCBE-DIDS have been deposited to the Electron Microscopy Data Bank under the accession numbers EMD-66593 (human NCBE with NaHCO_3_), EMD-66591 (human NCBE with KHCO_3_), EMD-66590 (human NCBE with NaCl) and EMD-66592 (human NCBE with DIDS), respectively. The coordinates of human NCBE-NaHCO_3_, NCBE-KHCO_3_, NCBE-NaCl and NCBE-DIDS have been deposited to the Protein Data Bank under the accession codes 9X5P, 9X5N, 9X5M and 9X5O, respectively. The code and associated files for the MD simulations are available on GitHub at https://github.com/Hanting-lab/MD-simulation/tree/main/XXX.

## Acknowledgments

We thank Yuli Jiang, Yi Lv, and Songjie Lv from the Institute for Translational Brain Research at Fudan University for their support with imaging and animal experiments. We thank Z. Chen and H. Zhao at the Center of cryo-EM at Fudan University for their support with cryo-EM data collection. This work was funded by the grants from the Ministry of Science and Technology of the People’s Republic of China (STI2030-Major Projects 2022ZD0212600, H.Y.; 2022YFA1106400 to Y.C.); National Natural Science Foundation of China (32171216, H.Y.; 82273304/32200640 to Y.C.; 82271499, J. D.; 82403445 to Y. H.; 82573334 to G. S.); Shanghai Academic Research Leader (22XD1402700, 998 to G. S.), Shanghai Health Commission Clinical Research Special Project 999 (202440012, to G. S.); China Postdoctoral Science Foundation (2023M730689, X.S.).

## Author contributions

H.Y. and P.S. conceived and designed the project. P.S. performed the single-cell RNA-seq data analysis and MD simulations. P.S., and X.W. purified the proteins for cryo-EM and biochemical analysis. Y.H. performed immunohistochemistry and immunoblotting of mouse tissue. P.S., X.S., and A. D. collected cryo-EM data. P.S. processed cryo-EM data, built and refined models with the help of J.K.Z., and analysed the structures under the supervision of H.Y. P.S. and X.W. performed the cell-based pH recovery assays, analysed and interpreted the results with the assistance of X.G. P.S., X.W., and H.Y. prepared the figures and wrote the manuscript with input from Y.H., J.K.Z., X.S., J.D., and Y.C.

## Corresponding author

Correspondence to

