## Supplemental figs and tables for "Substrate Coupling and Inhibition of the Human Na^+^-Dependent Cl^−^/HCO₃^−^ Exchanger"

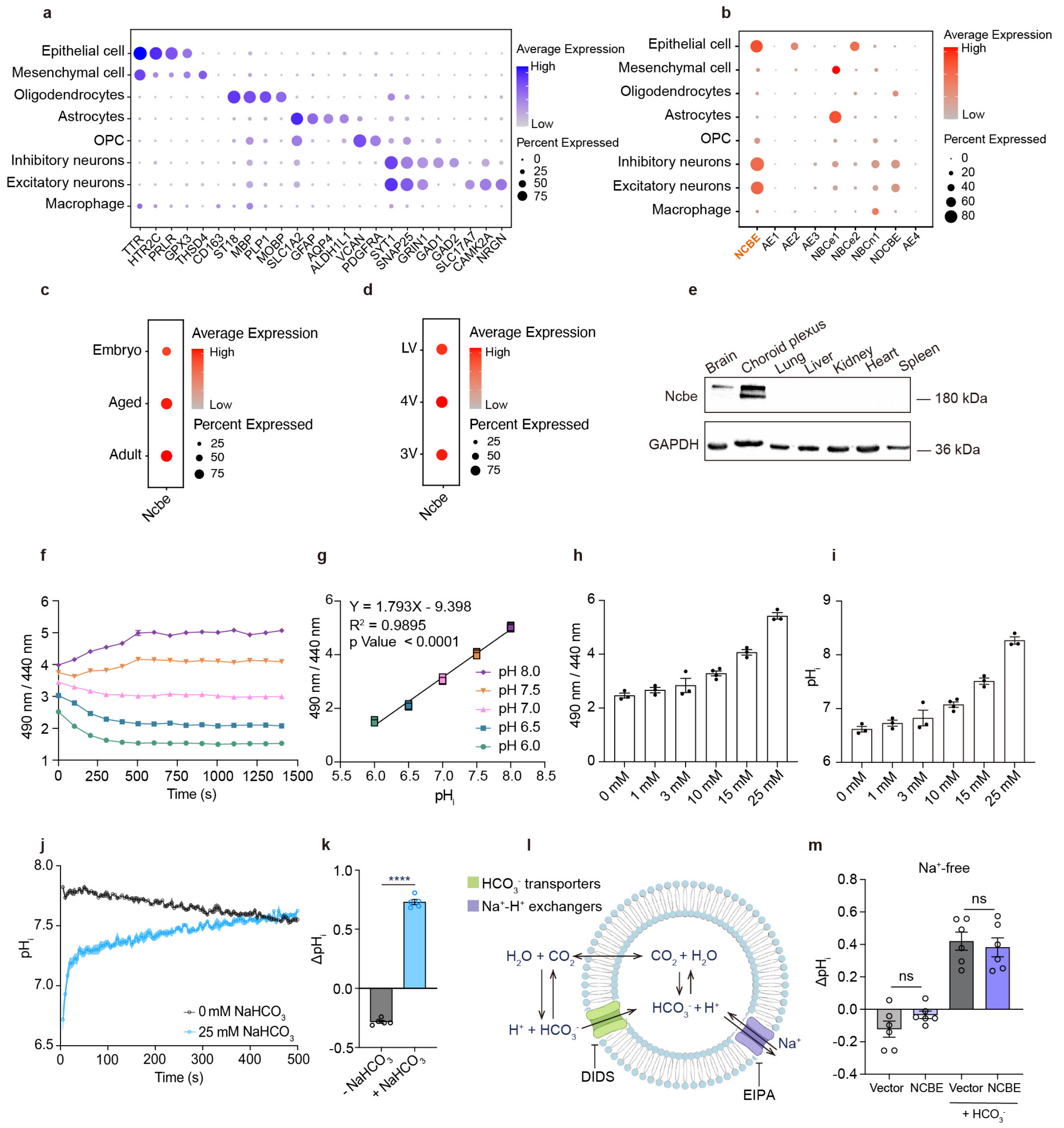


**Supplementary Fig 1. Functional characterization of NCBE, related to Figure 1.**

**a, b** Bubble plot presenting the cell-type-specific genes across the types of cells in human cortex and choroid plexus (a) and the SLC4 family transporters expression in different cell types (b). Original data were obtained from GEO: GSE159812. **c, d** Ncbe expression profiles in CP epithelial cells of embryonic, adult and aged mouse (c), and across lateral (LV), third (3V) and fourth (4V) ventricle (d). Original data were obtained from GEO: GSE168704. **e** Expression of Ncbe protein in brain, choroid plexus, lung, liver, kidney, heart and spleen of mouse. **f, g** Time course of the excitation fluorescence ratio (490 nm / 440 nm) (f) and calibration of the excitation fluorescence ratio into intracellular pH ($pH_i$) values (g) in HEK293T cells under high-$K^+$ treatment across different extracellular pH conditions, following exposure to a 10 μM mixture of nigericin and valinomycin. The experiments were performed in $n = 5$ biological replicates with each in technical triplicate. **h, i** Excitation fluorescence ratio (490nm / 440 nm) (h) and $pH_i$ (i) values in the presence of 0 mM, 1 mM, 3 mM, 10 mM, 15 mM and 25 mM $NaHCO_3$ for 10 min in HEK293T cells ($n$ =3). The experiments were performed in $n = 3$ technical triplicates. **j, k** Time course of $pH_i$ recovery (j) and $\Delta pH_i$ (k) in the presence of 25 mM $NaHCO_3$. $\Delta pH_i$ was calculated as the difference between pH at 0 s and 500 s. The experiments were performed in $n = 5$ biological replicates with each in technical triplicate. **l** Schematic depicts $HCO_3^-$-dependent $pH_i$ regulation mechanism of living cells. Intracellular pH recovery is governed primarily by two mechanisms. One involves $HCO_3^-$ transporters, which import bicarbonate into the cell, altering the $CO_2/HCO_3^-$ equilibrium and buffering intracellular protons. The other system relies on $Na^+/H^+$ exchangers that use the inward $Na^+$ gradient to extrude $H^+$, thereby elevating $pH_i$. EIPA blocks $Na^+/H^+$ exchangers and DIDS inhibits $HCO_3^-$ transporters. **m** $HCO_3^-$-dependent $\Delta pH_i$ under $Na^+$-free treatment in HEK293T cells ($n = 6$ biological replicates) related to Fig.1D. $\Delta pH_i$ was calculated as the difference between pH at 400 s and 690 s after the addition of acid-loading with $NH_4Cl$. The experiments were performed in technical triplicate. Data are presented as mean ± SEM. Statistical analysis was performed using unpaired t-test; ****, $P < 0.0001$, ns, $P \geq 0.05$.

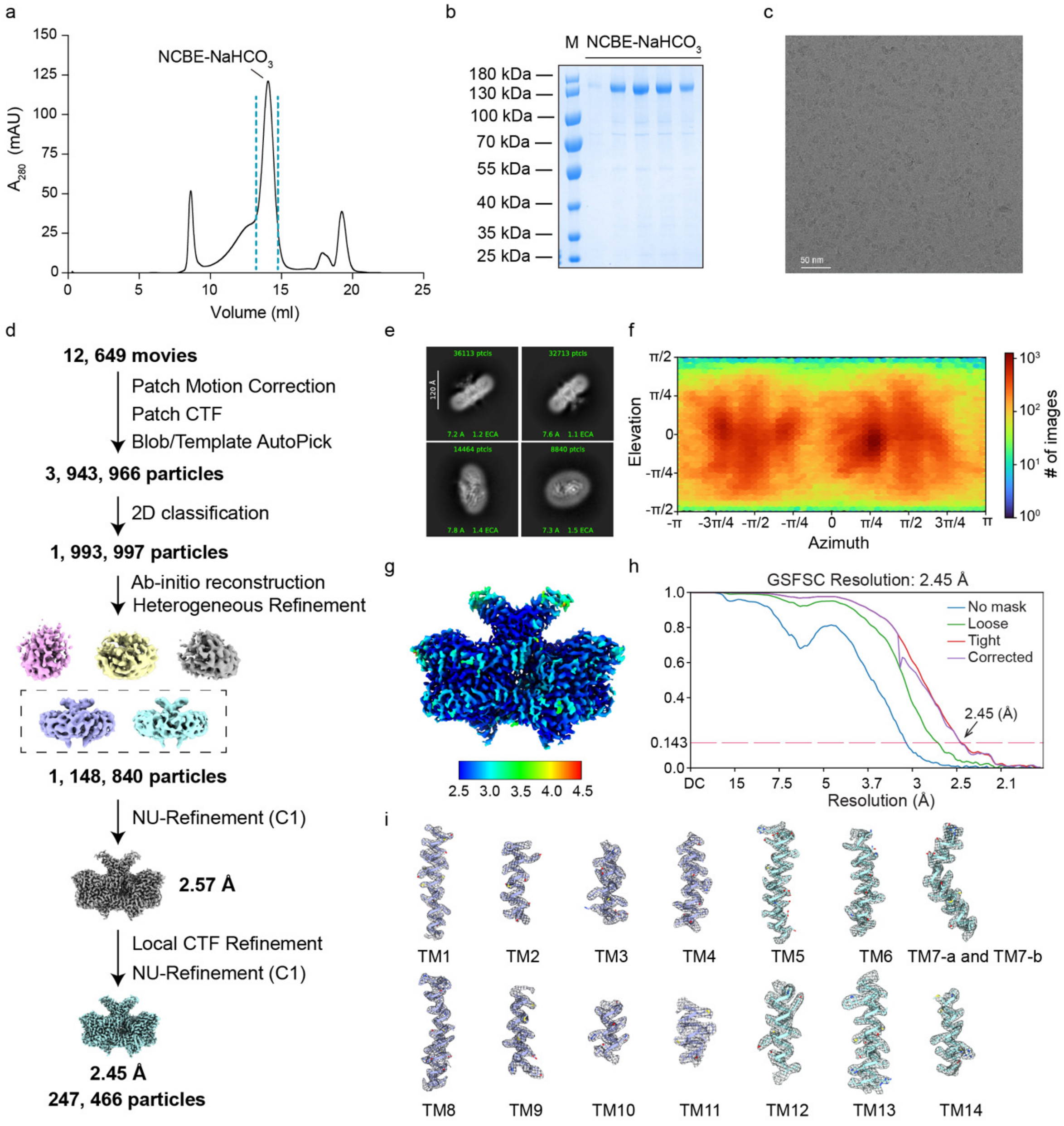


**Supplementary Fig 2. Cryo-EM data processing of human NCBE with $NaHCO_3$.**

**a-c** Representative trace of size-exclusion chromatography (a), SDS-PAGE (b), and cryo-EM micrographs (c) of purified human NCBE protein with $NaHCO_3$. **d** Workflow of image processing strategy. **e** Representative 2D class averages. **f** Particle orientation distributions of particles used for the final 3D reconstructions. **g** Local resolution map. **h** Gold-standard Fourier Shell Correlation curves. **i** Representative Cryo-EM densities for TM helices.

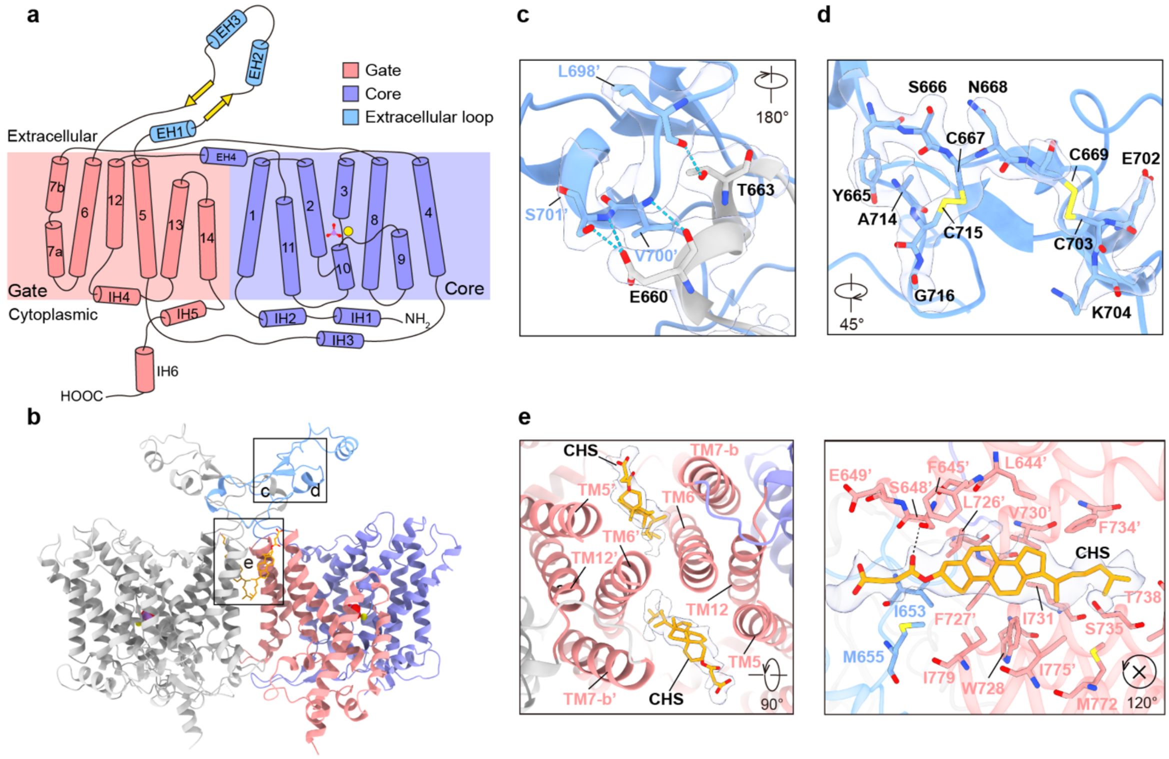


**Supplementary Fig 3. Hallmark features of homodimer NCBE structure with $NaHCO_3$.**

**a** TMD topology of human NCBE monomer. The core domain, gate domain, and extracellular loop are coloured purple, red, and blue, respectively. **b** The dimerization of NCBE involves the core and gate domains from the side view. The protomer of NCBE is segmented into three distinct regions: the core domain in purple, the gate domain in red, and the extracellular loop in blue. **c, d** Zoom-in views of the detailed interactions within the extracellular loop. Critical residues involved in inter-molecular (c) and intra-molecular (d) interactions are displayed as stick models and labelled. The density of the critical residues is shown in the transparency surface. **e** Zoomed in on the top (left) and side (right) views of the interaction between the core domain and CHS. CHS and critical residues are shown as sticks. The density of the CHS zoomed-in view is shown in a transparency surface.

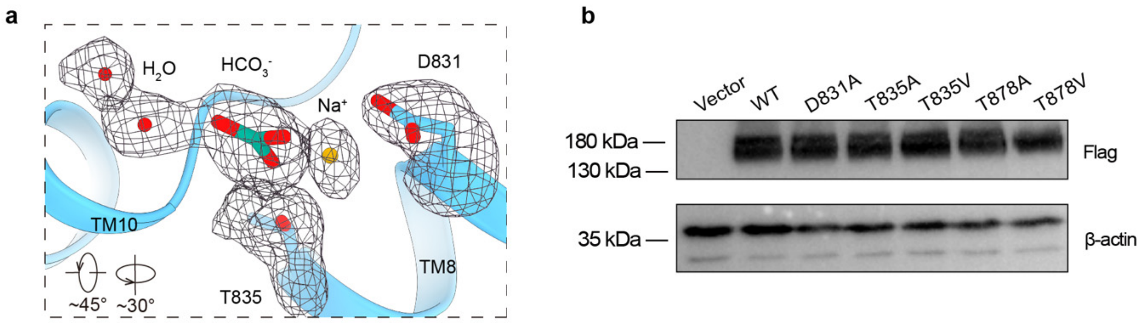


**Supplementary Fig 4. $Na^+$ and $HCO_3^-$ binding site and NCBE-mutants expression in HEK 293T cells, related to Fig. 2.**

**a** Zoom-in view of cryo-EM density for $Na^+$, $HCO_3^-$ and nearby residues in NCBE $NaHCO_3$-bound state within the conserved substrate-binding pocket. The density of ions, water and residues zoomed-in view was shown in black meshes at 0.125 level thresholds. Related to Fig. 2b from different view. **b** Immunoblotting analysis of NCBE and NCBE-mutants with Flag tag. Vector serves as control.

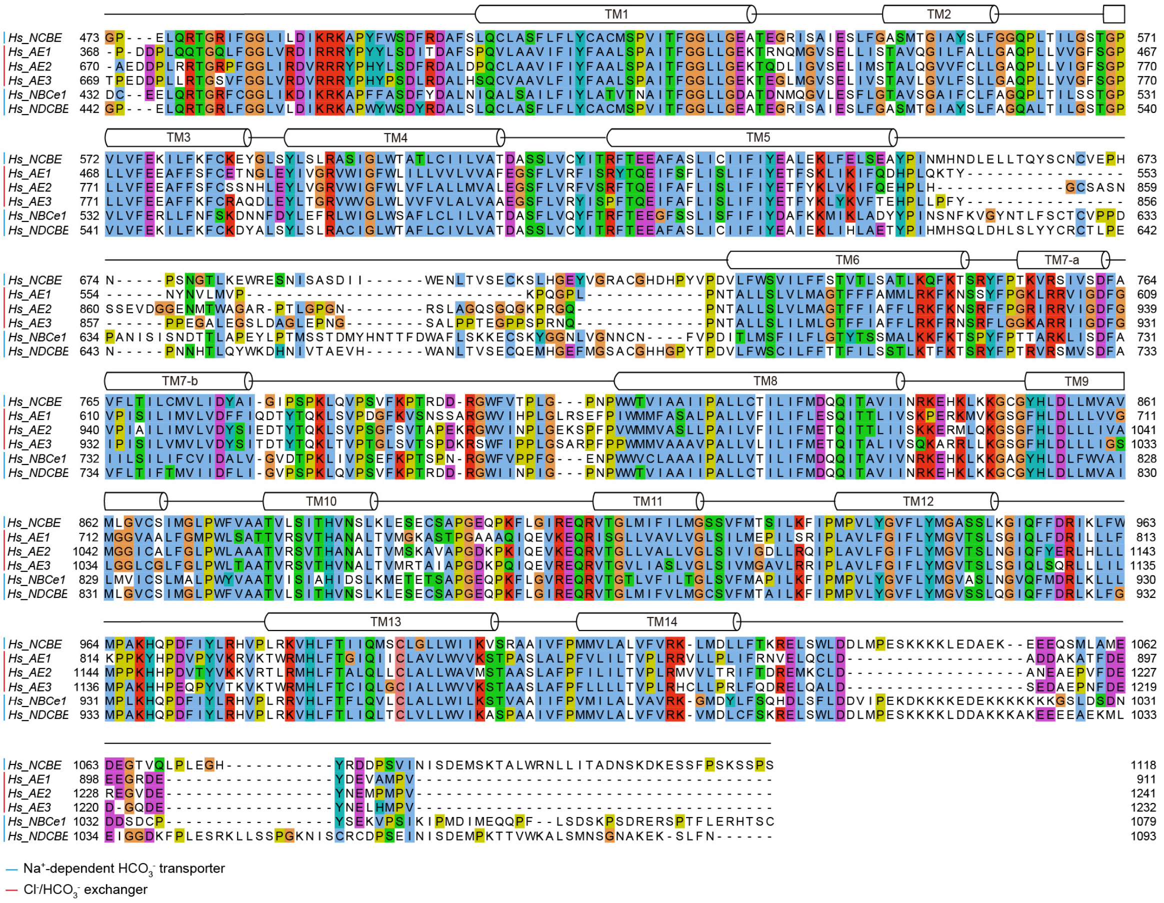


**Supplementary Fig 5. Sequence alignment of TM domains of the human SLC4 family.** Sequence alignments of TM domains of NCBE, AE1-3, NBCe1, and NDCBE from human using ClustalW software. Secondary structural elements are labelled on the top of the sequences.

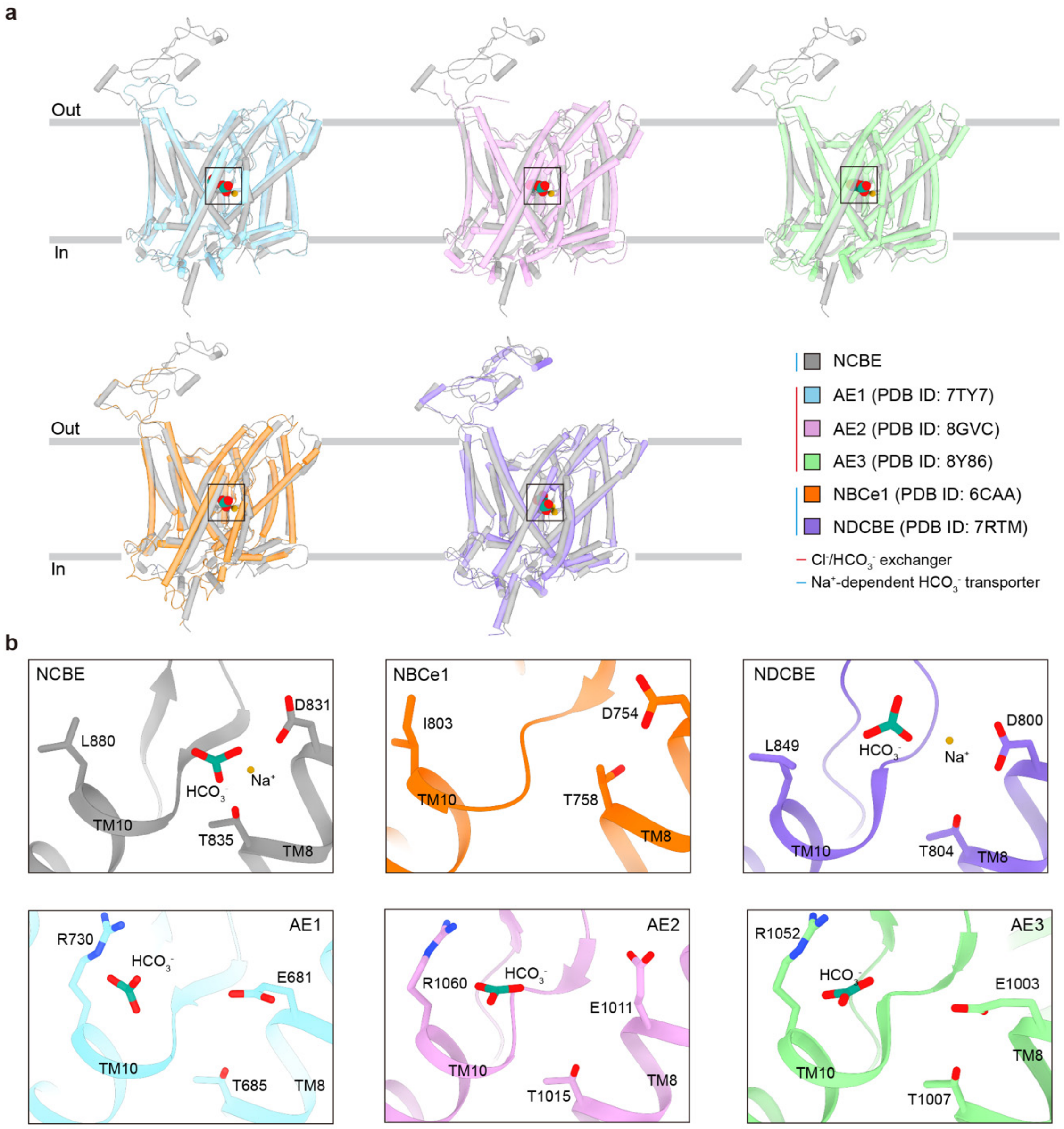


**Supplementary Fig 6. $HCO_3^-$-binding pockets of NCBE, AE1-3, NBCe1, and NDCBE**

**a** Structural superposition of the monomer structures of NCBE, AE1-3, NBCe1, and NDCBE. The proteins are coloured in grey, blue, pink, green, orange, and purple, respectively. **b** Zoomed-in views of the $HCO_3^-$-binding pockets of NCBE, AE1-3, NBCe1, and NDCBE. The proteins are coloured according to the same scheme as shown in (a). $Na^+$ and $HCO_3^-$ are represented as yellow and green sticks, respectively.

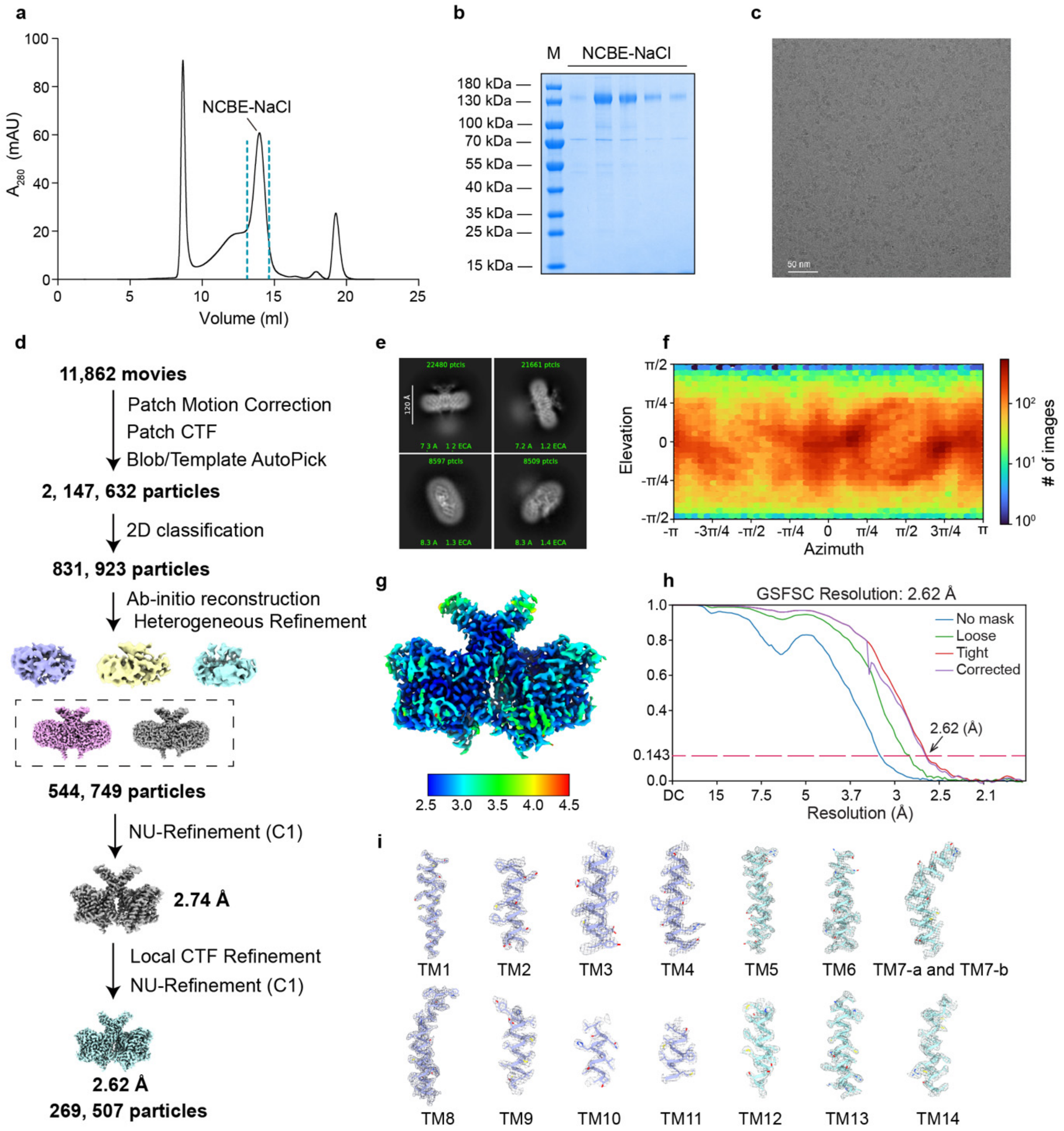


**Supplementary Fig 7. Cryo-EM data processing of human NCBE with NaCl.**

**a-c** Representative the trace of size-exclusion chromatography (a), SDS-PAGE (b), and cryo-EM micrographs (c) of purified human NCBE protein with NaCl. **d** Workflow of image processing strategy. **e** Representative 2D class averages. **f** Particle orientation distributions of particles used for the final 3D reconstructions. **g** Local resolution map. **h** Gold-standard Fourier Shell Correlation curves. **i** Representative Cryo-EM densities for TM helices.

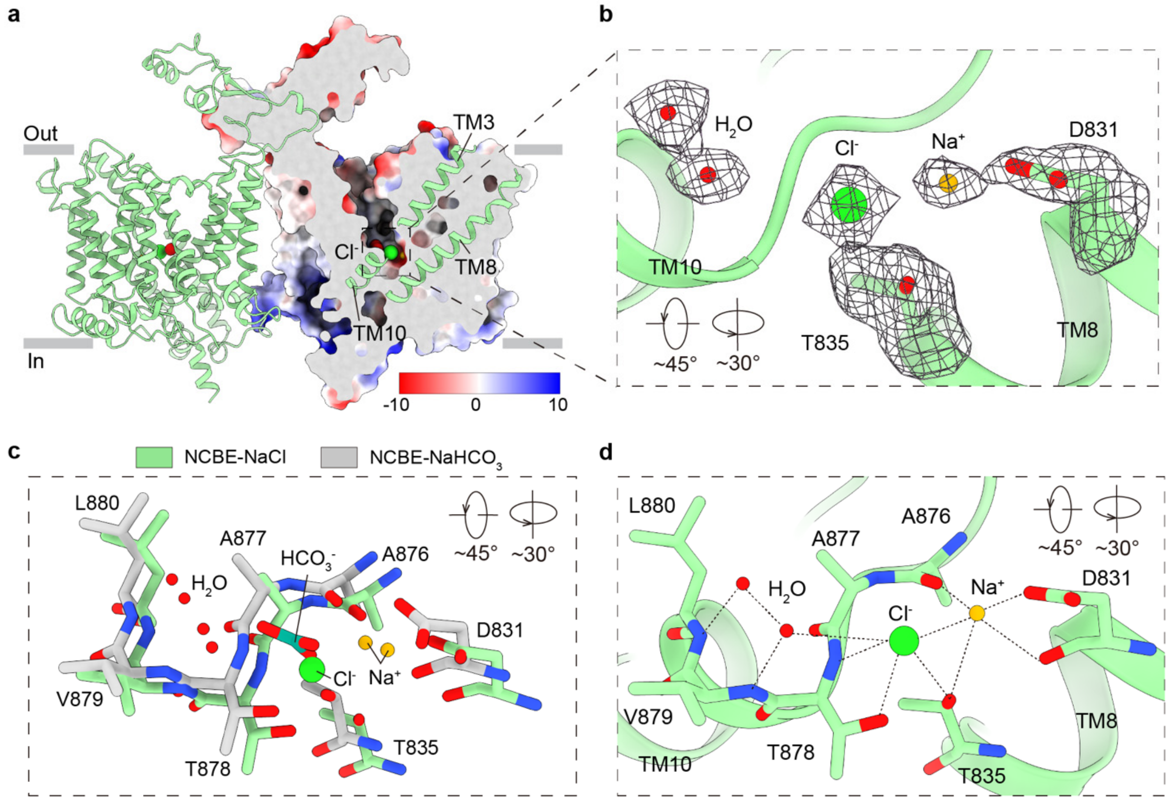


**Supplementary Fig 8. $Na^+$ and $Cl^-$ binding site of NCBE.**

**a** Ribbon model and electrostatic potential surface map of NBCE NaCl-bound state. The electrostatic surface map was sliced to show the $Cl^-$ ion bound (green sphere), with the helices TM3, TM8 and TM10 shown as green ribbon model. Regions with negative to positive charge are represented using a color gradient from red to blue. **b** Different view of Cryo-EM density for $Na^+$, $Cl^-$ and nearby residues in NCBE NaCl-bound state within the conserved substrate-binding pocket. The density of ions, water and residues zoomed-in view was shown in black meshes at 0.125 level thresholds. Related to Fig. 2h from different view. **c** Comparison of ion-binding site from NCBE $NaHCO_3$-bound state (grey) and NCBE NaCl-bound state (green). Related to Fig. 2j from different view. **d** Detailed view of NaCl binding site with key residues and coordination distances. Residues coordinated with $H_2O$, $Na^+$, and $Cl^-$ are depicted as atom sticks model.

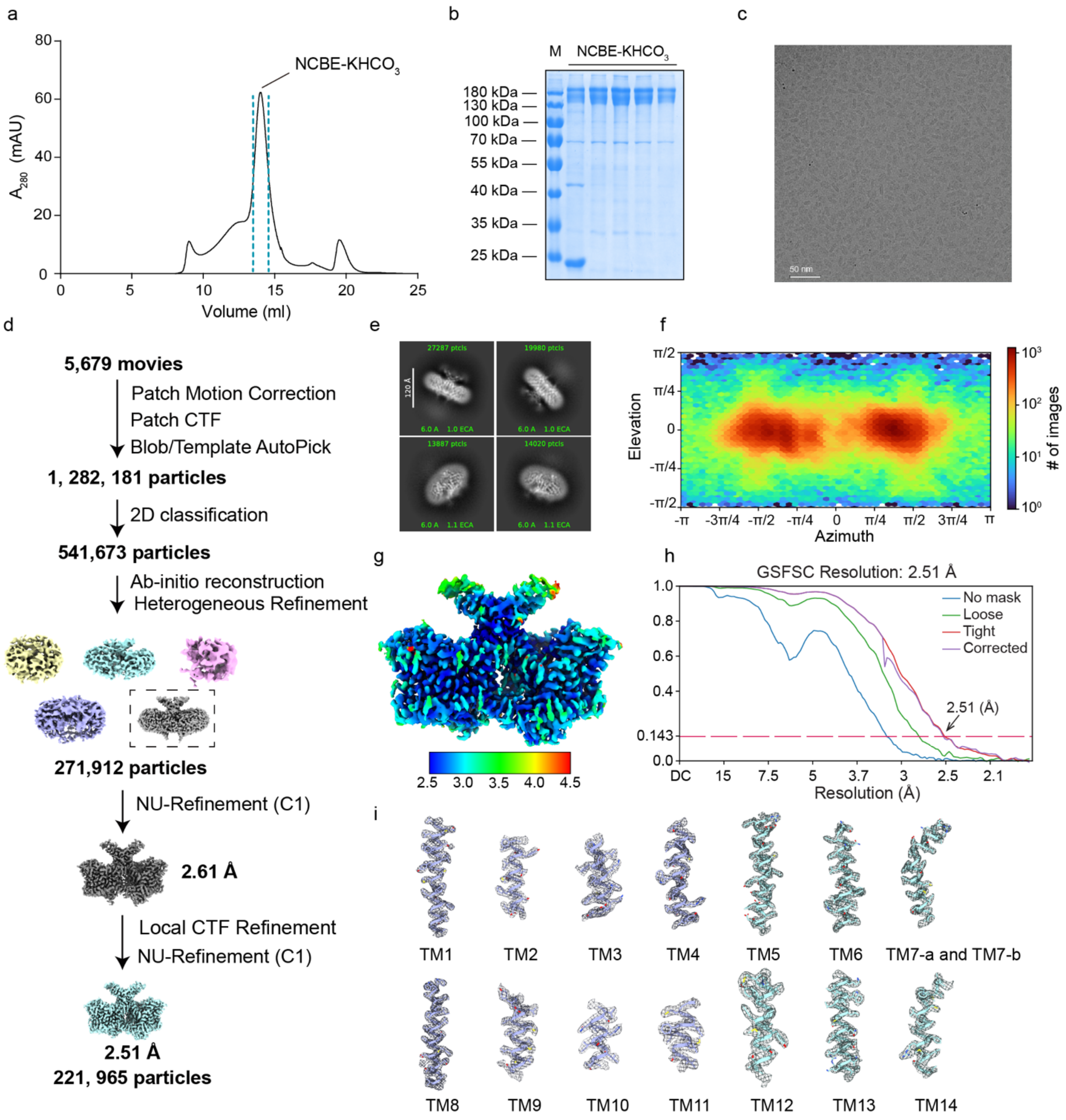


**Supplementary Fig 9. Cryo-EM data processing of human NCBE with $KHCO_3$.**

**a-c** Representative the trace of size-exclusion chromatography (a), SDS-PAGE (b), and cryo-EM micrographs (c) of purified human NCBE protein with $KHCO_3$. **d** Workflow of image processing strategy. **e** Representative 2D class averages. **f** Particle orientation distributions of particles used for the final 3D reconstructions. **g** Local resolution map. **h** Gold-standard Fourier Shell Correlation curves. **i** Representative Cryo-EM densities for TM helices.

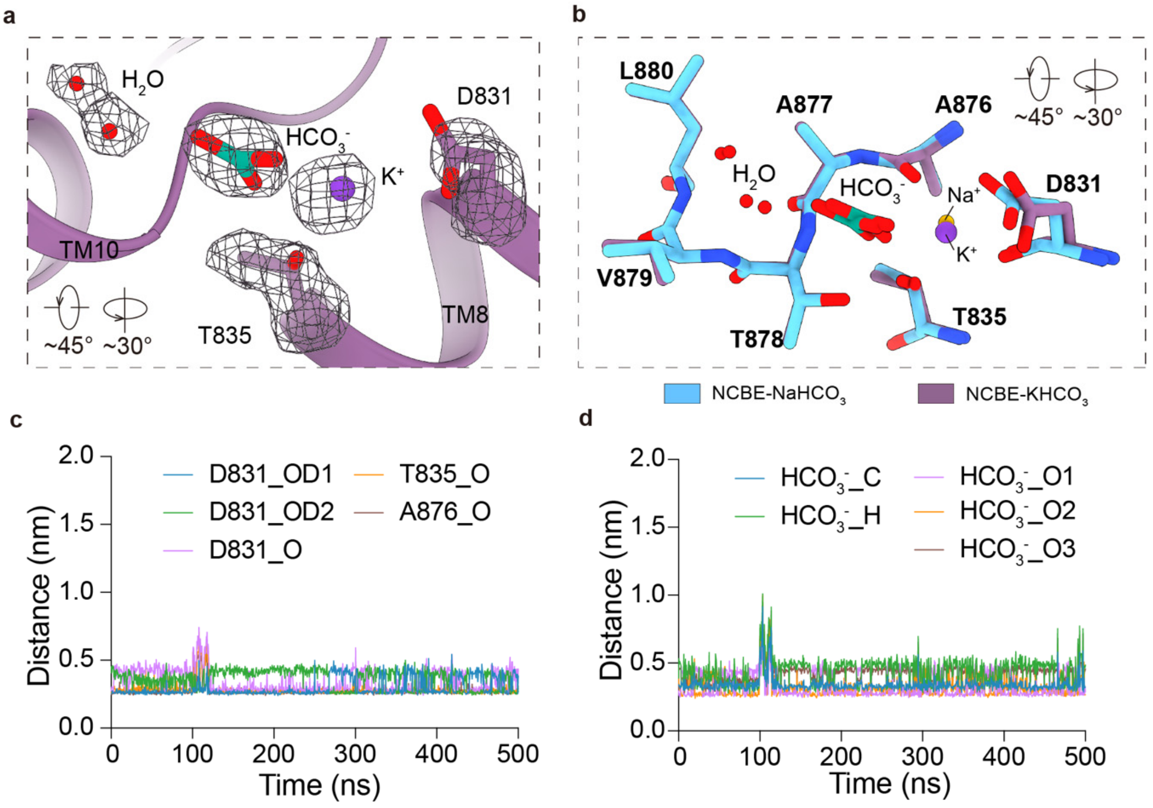


**Supplementary Fig 10. $K^+$ and $HCO_3^-$ binding site of NCBE, related to Fig. 3.**

**a** Different view of cryo-EM density for $K^+$, $HCO_3^-$ and nearby residues in NCBE $KHCO_3$-bound state within the conserved substrate-binding pocket. The density of ions, water and residues zoomed-in view was shown in black meshes at 0.125 level thresholds. Related to Fig. 3b from different view. **b** Comparison of ion-binding site from NCBE $KHCO_3$-bound state (purple) and NCBE $NaHCO_3$-bound state (blue). Related to Fig. 3c from different view. **c, d** MD simulation of 500 ns time scale illustrating the distance between the $K^+$ and its coordinated residues (c), as well as the distance between the $K^+$ and the $HCO_3^-$ (d).

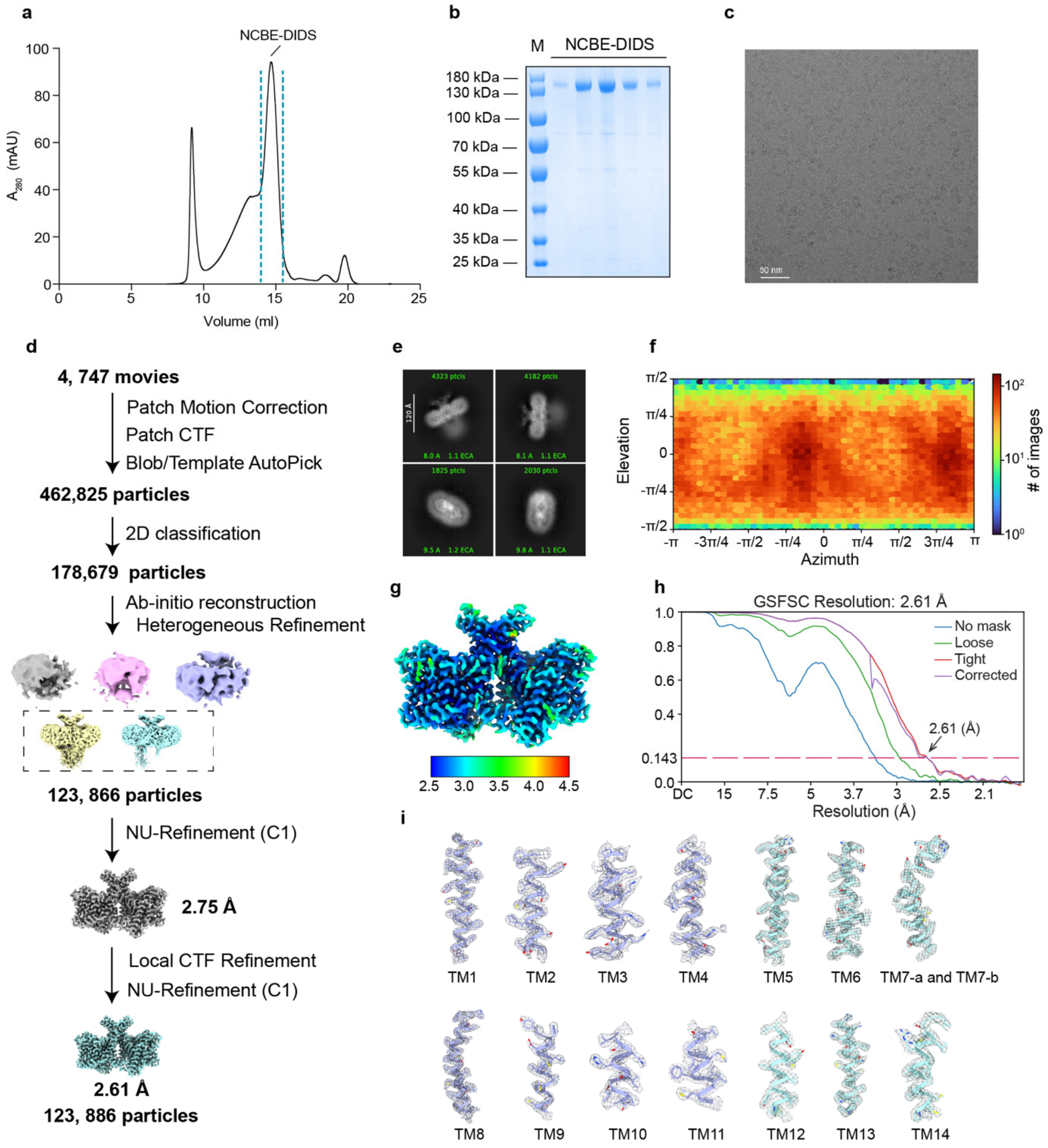


**Supplementary Fig 11. Cryo-EM data processing of human NCBE with DIDS.**

**a-c** Representative the trace of size-exclusion chromatography (a), SDS-PAGE (b), and cryo-EM micrographs (c) of purified human NCBE protein with DIDS. **d** Workflow of image processing strategy. **e** Representative 2D class averages. **f** Particle orientation distributions of particles used for the final 3D reconstructions. **g** Local resolution map. **h** Gold-standard Fourier Shell Correlation curves. **i** Representative Cryo-EM densities for TM helices.

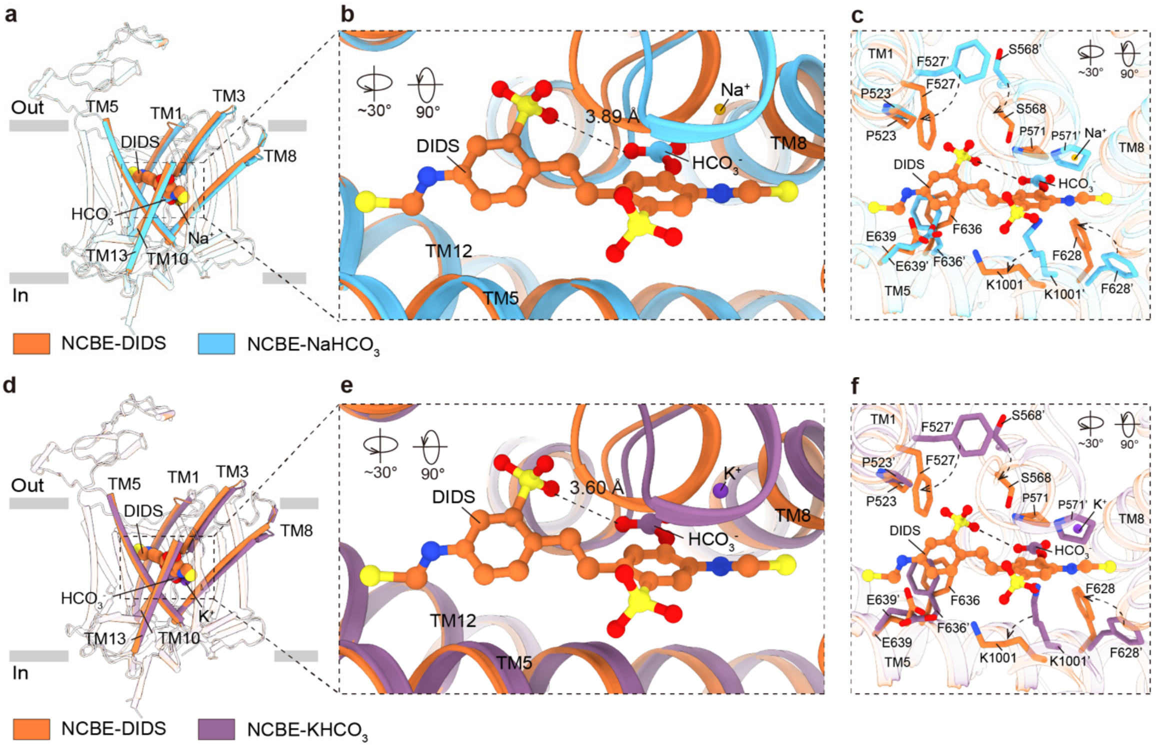


**Supplementary Fig 12. Structural comparison of NCBE-DIDS with NCBE-$NaHCO_3$ and NCBE-$KHCO_3$**

**a, d** Structural comparison of NCBE-DIDS (orange) with NCBE-$NaHCO_3$ (blue) (a) and NCBE-$KHCO_3$ (purple) (d). Critical TMs are represented as coloured cylinders without transparency. DIDS, $Na^+$, $K^+$ and $HCO_3^-$ are shown as sphere models. **b, e** Structural comparison of DIDS binding pocket with $NaHCO_3$ (b) and $KHCO_3$ (e) binding pocket. DIDS, $Na^+$, $K^{+,}$ and $HCO_3^-$ are shown as stick models. **c, f** Zoom-in view of the DIDS-binding pocket and interactions of the DIDS in the NCBE-DIDS structures. The DIDS is shown as an orange stick model. $Na^+$ and $HCO_3^-$ are shown as yellow and green stick models. Critical residues interacting with the ligands are shown as sticks and labelled.

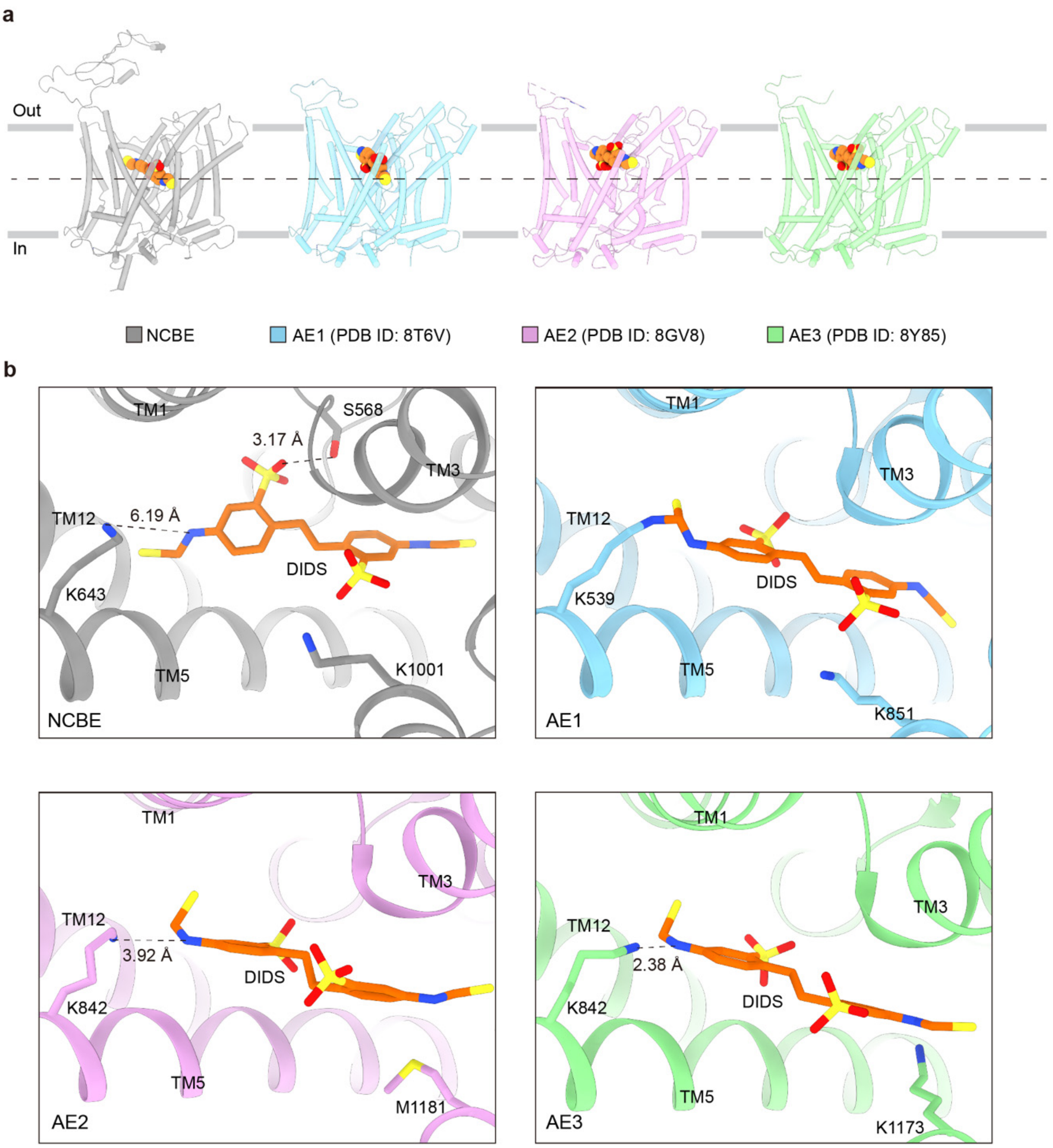


**Supplementary Fig 13. DIDS binding pockets of AE1-3 and NCBE.**

**a** Transparent cylinder cartoon representations of the AE1-3 and NCBE structures bound with DIDS. DIDS is shown as orange stick sphere model. NCBE and AE1-3 are colored by grey, blue, pink and green, respectively. **b** Zoom-in views of DIDS binding pockets of AE1-3 and NCBE. DIDS is shown as orange stick model. NCBE, AE1-3, NBCe1 and NDCBE are colored to the same scheme as show in (**a**).

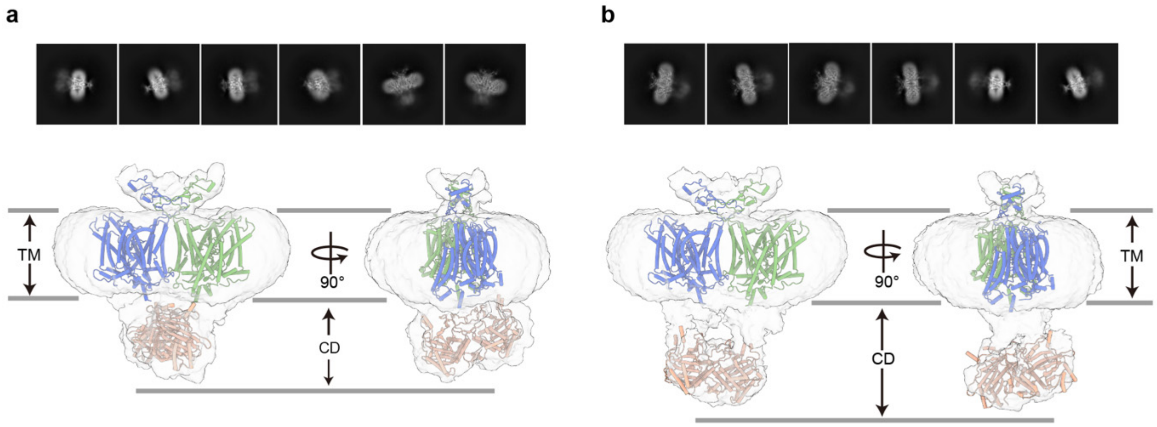


**Supplementary Fig 14. Visualization of cytoplasmic domain of NCBE bound with DIDS.**
**a** 2D projection and fitted model of the DIDS-bound state of NCBE with the CD positioned close to the transmembrane domain (TM). **b** 2D projection and fitted model of the DIDS-bound state of NCBE with the CD positioned away from the transmembrane domain (TM).

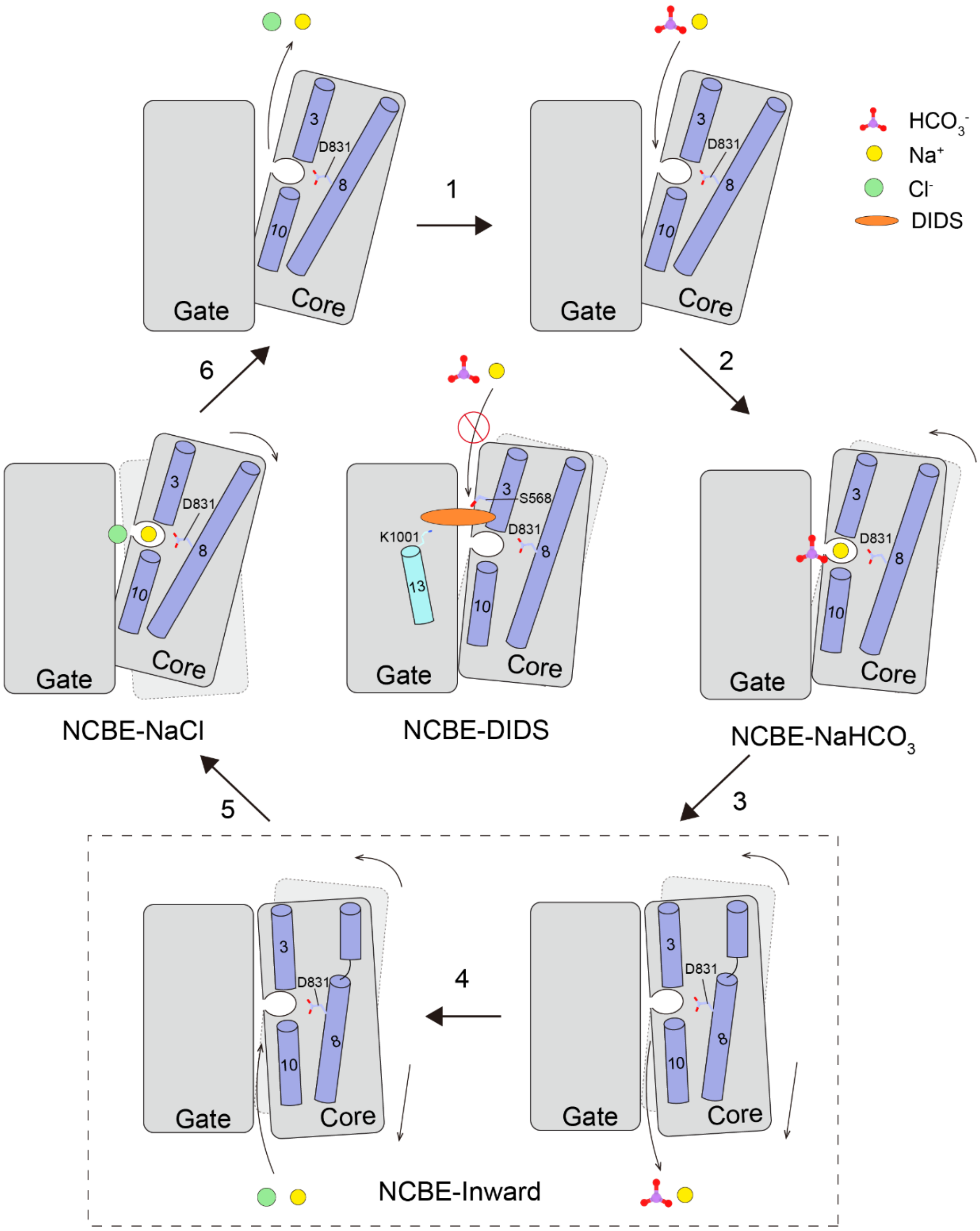


**Supplementary Fig 15. Proposed model of $Na^+$-dependent $Cl^-$/$HCO_3^-$ exchange mediated by NCBE and its inhibition by DIDS.**

Schematic depicts $Na^+$-dependent $Cl^-$-$HCO_3^-$ exchange mediated by NCBE and its inhibition by DIDS. Critical TM domains are highlighted in purple (TM3, TM8 and TM10) and cyan (TM13). Key residues are shown as stick model (S568, D831 and K1001). Proposed model suggests an elevator-like transport mechanism of NCBE.

**Table S1.**

**Cryo-EM data collection, refinement and validation statistics.**

| | NCBE-$NaHCO_3$ (EMDB-66593) (PDB 9X5P) | NCBE-NaCl (EMDB-66590) (PDB 9X5M) | NCBE-$KHCO_3$ (EMDB-66591) (PDB 9X5N) | NCBE-DIDS (EMDB-66592) (PDB 9X5O) |
|---|---|---|---|---|
| **Data collection and processing** | | | | |
| Magnification | 130,000x | 130,000x | 130,000x | 130,000x |
| Voltage (kV) | 300 | 300 | 300 | 300 |
| Electron exposure (e–/Å$^2$) | 50.17 | 49.10 | 53.93 | 52.58 |
| Defocus range (μm) | -0.8 to -1.8 | -0.8 to -1.8 | -0.8 to -1.8 | -0.8 to -1.8 |
| Pixel size (Å) | 0.932 | 0.932 | 0.932 | 0.932 |
| Symmetry imposed | C1 | C1 | C1 | C1 |
| Initial particle images (no.) | 3,943,966 | 2,147,632 | 1,282,181 | 560,247 |
| Final particle images (no.) | 624,534 | 269,507 | 221,965 | 123,886 |
| Map resolution (Å) | 2.39 | 2.62 | 2.51 | 2.61 |
| FSC threshold | 0.143 | 0.143 | 0.143 | 0.143 |
| Map resolution range (Å) | 2.072-3.014 | 2.338-44.442 | 2.114-39.561 | 2.372-44.169 |
| **Refinement** | | | | |
| Initial model used (PDB code) | AlphaFold3 | AlphaFold3 | AlphaFold3 | AlphaFold3 |
| Model resolution (Å) | 2.52 | 2.73 | 2.7 | 2.9 |
| FSC threshold | 0.5 | 0.5 | 0.5 | 0.5 |
| Model resolution range (Å) | NA | NA | NA | NA |
| Map sharpening *B* factor (Å$^2$) | -76 | -76 | -60 | -61.4 |
| Model composition | | | | |
| Non-hydrogen atoms | 9275 | 9159 | 9226 | 9188 |
| Protein residues | 1161 | 1146 | 1152 | 1142 |
| Ligands | NA:2 Y01:2 BCT:2 Water:4 | NA:2 Y01:2 Cl:2 Water: 4 | K:2 Y01:2 BCT:2 Water:4 | 4KU:2 Y01:2 |
| *B* factors (Å$^2$) | | | | |
| Protein | 36.97 | 39.99 | 64.85 | 57.46 |
| Ligand | 42.54 | 47.78 | 81.13 | 61.06 |
| R.m.s. deviations | | | | |
| Bond lengths (Å) | 0.003 | 0.002 | 0.003 | 0.003 |
| Bond angles (°) | 0.498 | 0.470 | 0.483 | 0.555 |
| Validation | | | | |
| MolProbity score | 1.61 | 1.41 | 1.72 | 1.91 |
| Clashscore | 5.79 | 2.37 | 6.20 | 6.70 |
| Poor rotamers (%) | 2.45 | 3.28 | 3.16 | 2.48 |
| Ramachandran plot | | | | |
| Favored (%) | 98.10 | 98.07 | 98.25 | 96.38 |
| Allowed (%) | 1.90 | 1.93 | 1.75 | 3.62 |
| Disallowed (%) | 0.00 | 0.00 | 0.00 | 0.00 |

**Table S2.**

**Molecular Dynamics simulation system for NCBE-$NaHCO_3$**

| **Molecule** | **Number of Molecules** |
|---|---|
| NCBE | 2 |
| POPC | 540 |
| $Na^+$ | 200 |
| $Cl^-$ | 197 |
| $HCO_3^-$ | 14 |
| Water | 67906 |
| Total atoms | 95185 |

**Table S3.**

**Molecular Dynamics simulation system for NCBE-NaCl**

| Molecule | Number of Molecules |
|---|---|
| NCBE | 2 |
| POPC | 535 |
| $Na^+$ | 191 |
| $Cl^-$ | 205 |
| Water | 68860 |
| Total atoms | 97103 |

**Table S4.**

**Molecular Dynamics simulation system for NCBE-$KHCO_3$**

| Molecule | Number of Molecules |
| --- | --- |
| NCBE | 2 |
| POPC | 540 |
| $K^+$ | 189 |
| $Cl^-$ | 188 |
| $HCO_3^-$ | 14 |
| Water | 68063 |
| Total atoms | 95517 |

**Table S5.**

**Molecular Dynamics simulation system for NCBE-DIDS**

| Molecule | Number of Molecules |
|---|---|
| NCBE | 2 |
| DIDS | 2 |
| POPC | 541 |
| $Na^+$ | 190 |
| $Cl^-$ | 189 |
| $HCO_3^-$ | 11 |
| Water | 65241 |
| Total atoms | 87167 |
